# The RNA-binding protein RBP42 regulates cellular energy metabolism in mammalian-infective *Trypanosoma brucei*

**DOI:** 10.1101/2021.04.14.439849

**Authors:** Anish Das, Tong Liu, Hong Li, Seema Husain

## Abstract

RNA-binding proteins are key players in coordinated post-transcriptional regulation of functionally related genes, defined as RNA regulons. RNA regulons play particularly critical roles in parasitic trypanosomes, which exhibit unregulated co-transcription of long arrays of unrelated genes. In this report, we present a systematic analysis of an essential RNA-binding protein, RBP42, in the mammalian-infective slender bloodstream form of African trypanosome, and we show that RBP42 is a key regulator of parasite’s central carbon and energy metabolism. Using individual-nucleotide resolution UV cross-linking and immunoprecipitation (iCLIP) to identify genome-wide RBP42-RNA interactions, we show that RBP42 preferentially binds within the coding region of mRNAs encoding core metabolic enzymes. Using global quantitative transcriptomic and proteomic analyses, we also show that loss of RBP42 reduces the abundance of target mRNA-encoded proteins, but not target mRNA, suggesting a plausible role of RBP42 as a positive regulator of target mRNA translation. Analysis reveals significant changes in central carbon metabolic intermediates following loss of RBP42, further supporting its critical role in cellular energy metabolism.

## Introduction

RNA regulons that coordinately regulate the production of functionally related proteins are emerging as key regulatory modules in eukaryotes^1^. In contrast to the prokaryotic ‘operons’ in which functionally related gene clusters are co-transcribed and co-translated, RNA regulons co-regulate gene cohorts post-transcriptionally by dynamic RNA-protein and RNA-RNA interactions to modulate critical regulatory steps, including messenger RNA (mRNA) maturation, localization, translation, and decay^2^. Thus, RNA regulons allow rapid, yet precise response to both intra- and extra-cellular signals that trigger readjustment of entire biochemical pathways^3^. Accordingly, RNA regulons have been discovered in diverse cell systems performing crucial regulatory functions^4,5^.

RNA regulon mediated post-transcriptional regulation is particularly important in *Trypanosoma brucei* subspecies, a deadly group of parasitic protozoa that cause Human African trypanosomiasis (HAT) and the cattle disease Nagana^6-8^. *T. brucei*, with a complex life cycle in two very different host environments, mammals and tsetse flies, undergo major morphological and metabolic changes that require precise coordination of specific gene expression patterns^9^. However, the profoundly unusual gene arrangement in *T. brucei*, where almost all protein-coding genes are co-transcribed as long polygenic arrays, prohibits the typical transcriptional regulation of individual genes^10,11^. Instead, regulation is primarily achieved at the post-transcriptional level^12^. Thus, RNA-binding proteins (RBPs), by interacting with specific sets of mRNAs, play key roles in controlling gene cohorts that are essential to maintain, or drive, developmental progression^7,13-14^. Recent studies have established critical roles of RBPs in trypanosome biology; overexpression of single RBP can force lifecycle-specific changes in the parasite^15,16^.

*T. brucei* central carbon and energy metabolism is a highly regulated process and undergoes striking changes in different host environments, dictated primarily by the availability of nutrients^17,18^. For example, in the tsetse fly midgut, the procyclic form harbors full repertoire of glycolytic enzymes as well as mitochondrial Krebs cycle and electron transport chain^19^. Although in glucose-rich culture media the procyclic forms preferentially metabolize glucose through glycolysis, in the fly midgut, abundant amino acids, such as proline is metabolized in the mitochondria and ATP is produced mainly by oxidative phosphorylation via a canonical cytochrome-containing electron transport chain^20^. In contrast, in mammalian blood, the slender bloodstream form uses glycolysis as the sole source of ATP^21,22^. Existing evidence suggest that slender bloodstream forms have no detectable Krebs cycle or mitochondrial respiratory chain coupled to ATP synthesis. Similar metabolic adjustments are also observed in trypanosomes growing in other varying environmental niches^23,24^. Although the molecular mechanisms that enable these dramatic metabolic remodeling are not yet fully understood, studies suggest critical roles performed by RBPs in this regulation^25-27^.

*T. brucei* RNA-binding protein RBP42 shares sequence homology to metazoan RAS-GTPase activating protein SH3 domain binding protein (G3BP)^28-30^. Metazoan G3BP proteins are a family of mRNA-binding protein that are known to regulate gene expression in response to intra- and extra-cellular stimuli. Multiple modes of function are attributed to G3BP proteins, which include mRNA stabilization and destabilization, subcellular localization and translation^31,32^. RBP42 is known to interact with translating, polysome-associated mRNAs encoding enzymatic proteins of central carbon metabolic pathways in procyclic trypanosomes^28^. However, the significance of these interactions in trypanosome gene regulation remain unknown.

In this report, we establish RBP42 as a post-transcriptional regulator of trypanosome central carbon and energy metabolism. Using protein-RNA crosslinking via ultraviolet (UV)-irradiation, we captured in vivo RBP42-RNA interactions in mammalian-infective slender bloodstream form *T. brucei*^33^. Identification of RBP42 crosslink sites on target transcripts shows that RBP42 binds within the coding sequence of many mRNAs that encode enzymatic proteins involved in cellular core metabolic pathways. Applying a conditional knockdown system, we show that RBP42 does not affect target-mRNA stability; loss of RBP42 has minimal effect on target mRNA abundance. Instead, RBP42 promotes target-mRNA translation; loss of RBP42 markedly reduces target mRNA-encoded protein abundance. Importantly, loss of RBP42 causes drastic reduction of many metabolic intermediates of the central carbon metabolic process, as well as ATP, NAD, and NADP that are critical for cellular energy and redox homeostasis. These data indicate that RBP42 plays an essential role in fine-tuning trypanosome energy metabolism, whose regulation is essential for *T. brucei*’s parasitic lifestyle.

## Results

### RBP42 binds within the coding region of transcripts encoding cellular primary metabolic enzymes

To accurately identify RBP42’s targets in mammalian-infective slender bloodstream form *T. brucei*, we performed UV cross-linking and immunoprecipitation (CLIP) by incorporating two strategies. First, to eliminate non-specific protein-antibody interactions from our analysis, we used a second independent antibody reagent, a mouse monoclonal antibody to Ty1-epitope (anti-Ty1), in addition to a rabbit polyclonal antibody to bacteria-derived recombinant RBP42 protein (anti-RBP42)^28^. Transgenic cell line (3Ty1-RBP42), producing triple-Ty1 epitope-tagged RBP42 protein, was generated by homologous recombination-mediated in situ tagging of both alleles of the diploid parasite (Figure 1a). Genomic PCR and immunoblot analyses of two representative clonal cell lines show proper integration and expression of the tagged-RBP42 protein (Figure 1b and c). 3Ty1-RBP42 cells that solely depend on tagged-RBP42 protein grow comparably to wild-type (WT) cells, demonstrating proper function of the tagged protein (Figure 1d). Both antibodies, anti-Ty1 and anti-RBP42, efficiently immunopurified crosslinked RBP42-RNA complexes from UV-irradiated 3Ty1-RBP42 cellular extracts. However, only anti-RBP42 efficiently immunopurified RBP42-RNA complexes from UV-irradiated WT cellular extracts, as expected (Figure 1e). We used WT cells in combination with anti-RBP42 antibody, and 3Ty1-RBP42 cells in combination with anti-Ty1 antibody, to identify RBP42’s cellular RNA targets (Figure 1e and f). Second, since UV-irradiated cross-linking of protein-RNA interactions are known to be very inefficient^34,35^, we captured RBP42-RNA interactions at two increasing UV doses, at 150mJ/cm^2^ and at 300mJ/cm^2^ constant energy. We reasoned that authentic RBP42-RNA interactions, as opposed to nonspecific, cryptic interactions, would be captured at both conditions.

**Fig 1.**
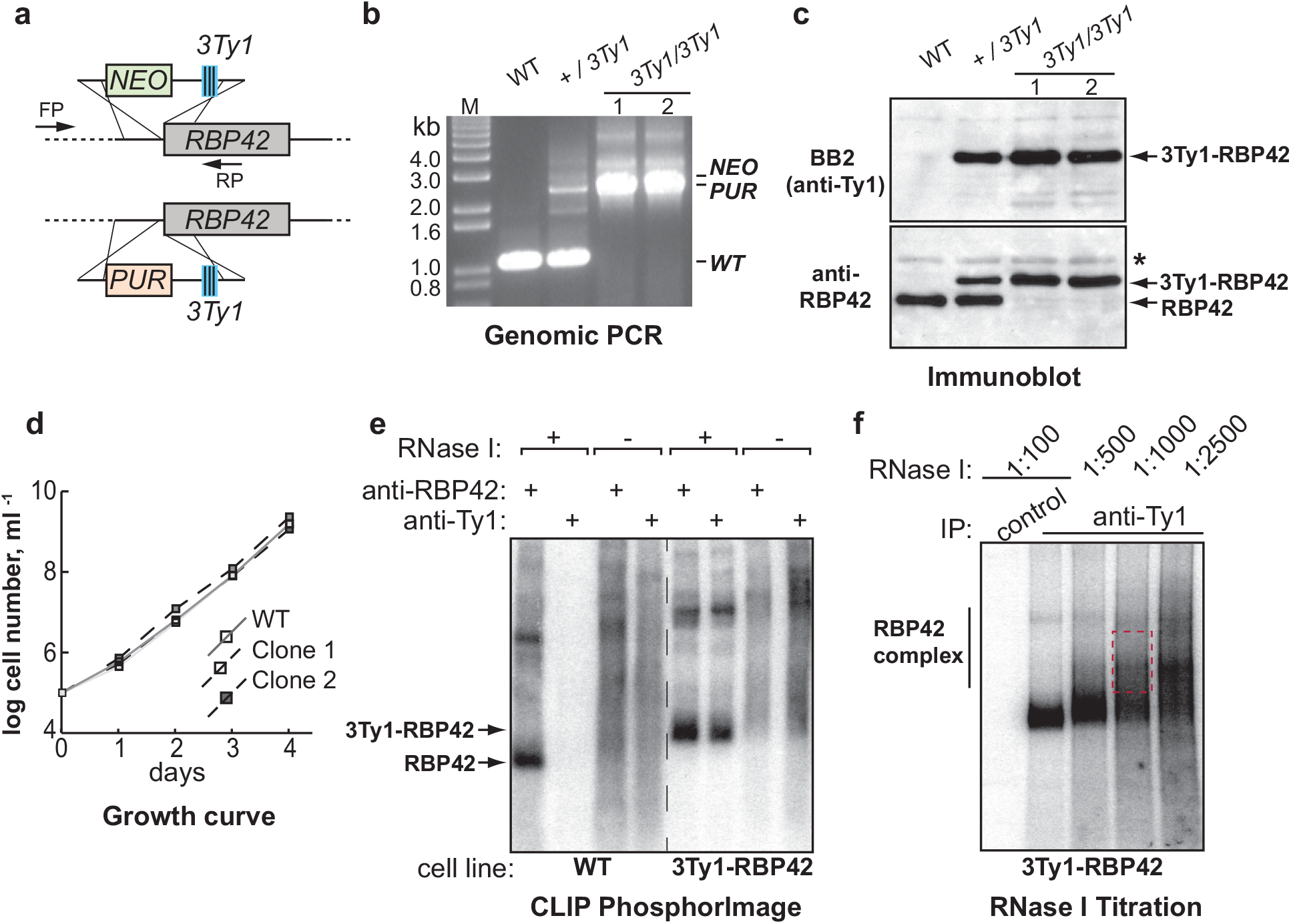
Crosslink Immunoprecipitation of RBP42-RNA complex from slender bloodstream form *T. brucei*. **(a)** Schematic showing homologous recombination-mediated in situ tagging to generate homozygous 3Ty1-epitope tagged RBP42 cell line. PCR generated DNA cassettes, containing neomycin (*NEO*) and puromycin (*PUR*) selectable marker genes, are shown. **(b)** Analysis of genomic DNA by PCR using FP and RP primers shown in panel (a) verifies proper integration of the DNA cassettes in two representative clonal cell lines. **(c)** Immunoblot analysis of total cellular proteins confirms the presence of only 3Ty1-tagged RBP42 protein in these clonal transgenic cell lines. Whereas anti-RBP42 antibody detects both native RBP42 and 3Ty1-tagged RBP42 proteins, anti-Ty1 (BB2) antibody detects only the tagged protein. Asterisk marks non-specific binding of anti-RBP42 antibody to an unknown protein. **(d)** Growth analysis shows that both clonal cell lines grow comparably to wild-type (WT) cells. **(e)** Immunoprecipitated protein-RNA complexes that were 5’-end radiolabeled on their RNA. Complexes were separated by denaturing polyacrylamide gel electrophoresis (SDS-PAGE) and transferred to nitrocellulose membranes. A PhosphorImage™ scan of ^32^P-labeled protein-RNA complexes purified from both wild-type (WT) cells and 3Ty1-RBP42 cells is shown. In the absence of ribonuclease (RNase I) the protein-RNA complexes are too large, which fail to resolve by gel chromatography, and appear as broad smear. While anti-Ty1 (BB2) antibody immunoprecipitated only the tagged protein, anti-RBP42 antibody immunoprecipitated both WT and tagged proteins. **(f)** Optimization of RNase I concentration to obtain optimal size RNA fragments for iCLIP library preparation. A representative PhosphorImager™ scan, similar to panel (e) is shown. Protein-RNA complexes were treated with decreasing concentrations of RNase I before immunoprecipitation. Complexes from the marked gel area (red dotted box) were excised from the nitrocellulose membrane, and extracted RNAs were reverse transcribed to prepare cDNA library. Normal mouse antibody serves as control.

We identified RBP42-RNA interactions by generating and deep-sequencing cDNA libraries from CLIP-purified RNA (Figure 1f). cDNA libraries were made using individual-nucleotide resolution CLIP (iCLIP) method^36^. The iCLIP, by incorporating a cDNA circularization step, takes advantage of the frequent termination of reverse transcriptase (RT) at the crosslink site, which therefore corresponds to the nucleotide preceding the cDNA start. Each unique cDNA molecule in the library denotes an individual crosslink event. We produced four independent iCLIP libraries: two for each of anti-RBP42 antibody (Samples 1 and 3) and anti-Ty1 antibody (Samples 2 and 4) at both 150mJ/cm^2^ (Samples 1 and 2) and 300mJ/cm^2^ UV doses (Samples 3 and 4) (Figure 2a). We obtained 29,464 and 26,055 unique cDNA molecules in Samples 1 and 2, respectively, and 867,214 and 493,361 unique cDNA molecules in Samples 3 and 4, respectively (Figure S1a). The large, ∼ 20-fold more unique cDNA molecules in Samples 3 and 4, compared to Samples 1 and 2 may occur as a batch effect, i.e., minor variations in enzymatic activities, and/or reaction conditions etc. It is also possible that higher, 300mJ/cm^2^, UV doses in Samples 3 and 4 produced more crosslink events (Figure S1b). Genome-wide comparison of mapped crosslink-sites reveals moderate to high degree of correlations among all four iCLIP libraries (Figure S1c). Distribution of crosslink-sites over annotated genomic features show that vast majority of RBP42 interactions occur within the mRNA coding sequences (Figure 2b). This result, in accordance with previously published data in fly-infective procyclic forms^28^, confirms that RBP42 predominantly binds within the target mRNA-coding sequences.

**Fig 2.**
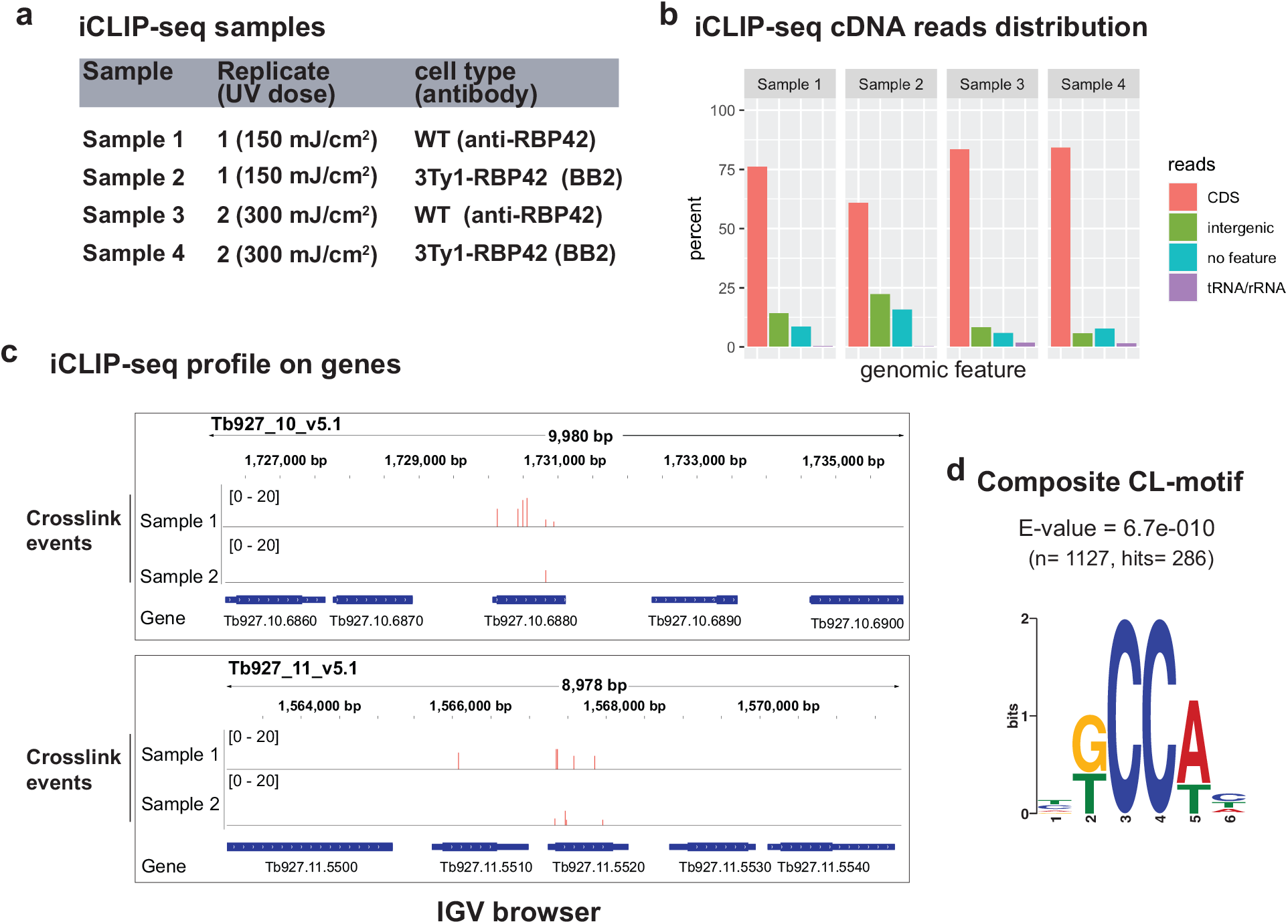
iCLIP-seq identifies RBP42’s RNA targets in slender bloodstream form *T. brucei*. **(a)** Four iCLIP-seq cDNA libraries (Samples 1 – 4) were made from both wild-type (WT) cells, using anti-RBP42 antibody (Samples 1 and 3), and 3Ty1-RBP42 cells using anti-Ty1 (BB2) antibody (Samples 2 and 4), applying two increasing UV doses, 150 mJ/cm^2^ (Sample 1 and 2) and 300 mJ/cm^2^ (Samples 3 and 4). **(b)** RBP42 predominantly binds within the coding region of mRNA targets. Bar plots of cDNA read distributions, mapped to *T. brucei* genomic features (*T. brucei brucei* strain 927 genome from TriTrypDB.org – version 46), are shown. CDS and intergenic denote the protein-coding open reading frames, and the 5’ and 3’ untranslated regions of mRNA respectively. tRNA/ rRNAs are as defined in the database mentioned above, and no features are without any annotation. **(c)** IGV browser view of RBP42 crosslinking to two candidate mRNA targets, one on chromosome 10 (Tb927.10.6880 glyceraldehyde 3-phosphate dehydrogenase, cytosolic), and the other on chromosome 11 (Tb927.11.5520 - triosephosphate isomerase). A crosslink event is counted for each unique cDNA and assigned to the upstream crosslink nucleotide (see method). The red bars indicate the number of crosslink events on crosslink sites. Few neighboring non-target genes are also shown as internal controls to indicate specificity of RBP42 interactions. **(d)** The composite sequence motif associated with RBP42 is discovered by comparing significant crosslink clusters using DREME algorithm (meme-suite.org).

We identified RBP42-target transcripts by recording crosslink-sites to annotated *T. brucei* 927 genome (TriTrypDB.org; V45). The number of crosslink events at each recorded crosslink-sites were calculated by counting number of stacked unique cDNAs with coinciding start sites (see method) (Figure 2c and S2). Evaluation by PureCLIP algorithm^37^, designed to analyze iCLIP data, identified similar crosslink sites. To identify potential RBP42 crosslink sequence motif, we sampled ∼1100 high-confidence crosslink site sequences, each 11 nucleotides (nt) long, consisting of crosslink site plus 5 nt flanking sequences, and searched for motif discovery using MEME suite (meme-suite.org/tools/meme). A consensus hexanucleotide sequence, centered around a CC dinucleotide, with highly significant enrichment (E = 6.7e-010) is revealed as the preferential RBP42 crosslink sequence motif (Figure 2d).

Tabulation of RBP42-targets, compiled from all four samples, resulted a combined and overlapping set of 2145 transcripts: 796 in Sample 1, 903 in Sample 2, 1449 in Sample 3, and 1020 in Sample 4 (Figure S1d). To undertake functional significance of RBP42 binding on target transcripts, we classified the ‘most reliable’ RBP42-target set using a stringent criterion – 189 congruent transcripts identified in all four iCLIP libraries. The 189 transcripts encode 181 ortholog protein groups that include 16 of unknown function and therefore annotated as hypothetical protein. Gene Ontology (GO) analysis of 165 known protein groups shows several enriched GO terms associated with primary metabolic processes (Figure S3). Similar result was reported with RBP42-targets in fly-infective procyclic forms^28^, indicating a conserved RBP42 function in both procyclic and slender bloodstream stages of the parasite. We also observed that RBP42-targets include transcripts encoding several RNA binding proteins, nine in the ‘most reliable’ 189-target set including its own, indicating a critical RBP42 function in modulating parasite’s RBP-mediated post-transcriptional regulation. Interestingly, many tRNAs, five tRNAs in the ‘most reliable’ set, were crosslinked to RBP42 (Supplementary Table). Since RBP42 is known to be associated with translating polysomes^28^, crosslinking to tRNAs is not surprising. However, the exact mechanism of how RBP42 connects both mRNA and tRNA within the translating polysomes is not known.

### RBP42 is essential for slender bloodstream form *T. brucei* survival

To investigate functional significance of RBP42 binding on target gene expression, we generated an RBP42 conditional knockdown slender bloodstream form cell line (RBP42^Ty1^) in which cell growth depends upon a tetracycline-regulated exogenously-expressed Ty1-tagged version of RBP42 (Figure 3a and d). Two antibiotic resistance genes replaced two native RBP42 alleles, which was confirmed by genomic PCR (Figure 3b). Immunoblot analysis confirmed that only the tagged RBP42 protein is expressed in RBP42^Ty1^ cells (Figure 3c). In the presence of tetracycline, added to the growth media, RBP42^Ty1^ cells grew normally. However, loss of RBP42, in the absence of tetracycline, caused RBP42^Ty1^ cells to stop dividing after two days, as reflected by the growth curve, and eventual death (Figure 3d). Immunoblot and immunofluorescence microscopic analyses confirmed the expected reduction of tagged-RBP42 protein in RBP42^Ty1^ cells (Figure 3e and S4a). RBP42 knockdown triggered marked phenotypic alterations, with cells exhibiting abnormal shape, and containing multiple nuclei (Figure S4a). Cell cycle analysis by flow cytometry reveals large increase in multinucleated (> 2) cells; ∼30% of cells following two days of RBP42 knockdown, compared to only ∼2% of normally growing cells, indicating apparent cytokinesis defects (Figure S4b). RBP42 knockdown triggered similar phenotypic alterations of procyclic forms ^28^, indicating its essential role in both mammalian and insect stages of the parasite. To seek answers to how RBP42 regulates *T. brucei* gene expression, we measured global changes in cellular transcriptome, proteome and metabolome in RBP42^Ty1^ cells following loss of RBP42 (Figure 3f).

**Fig 3.**
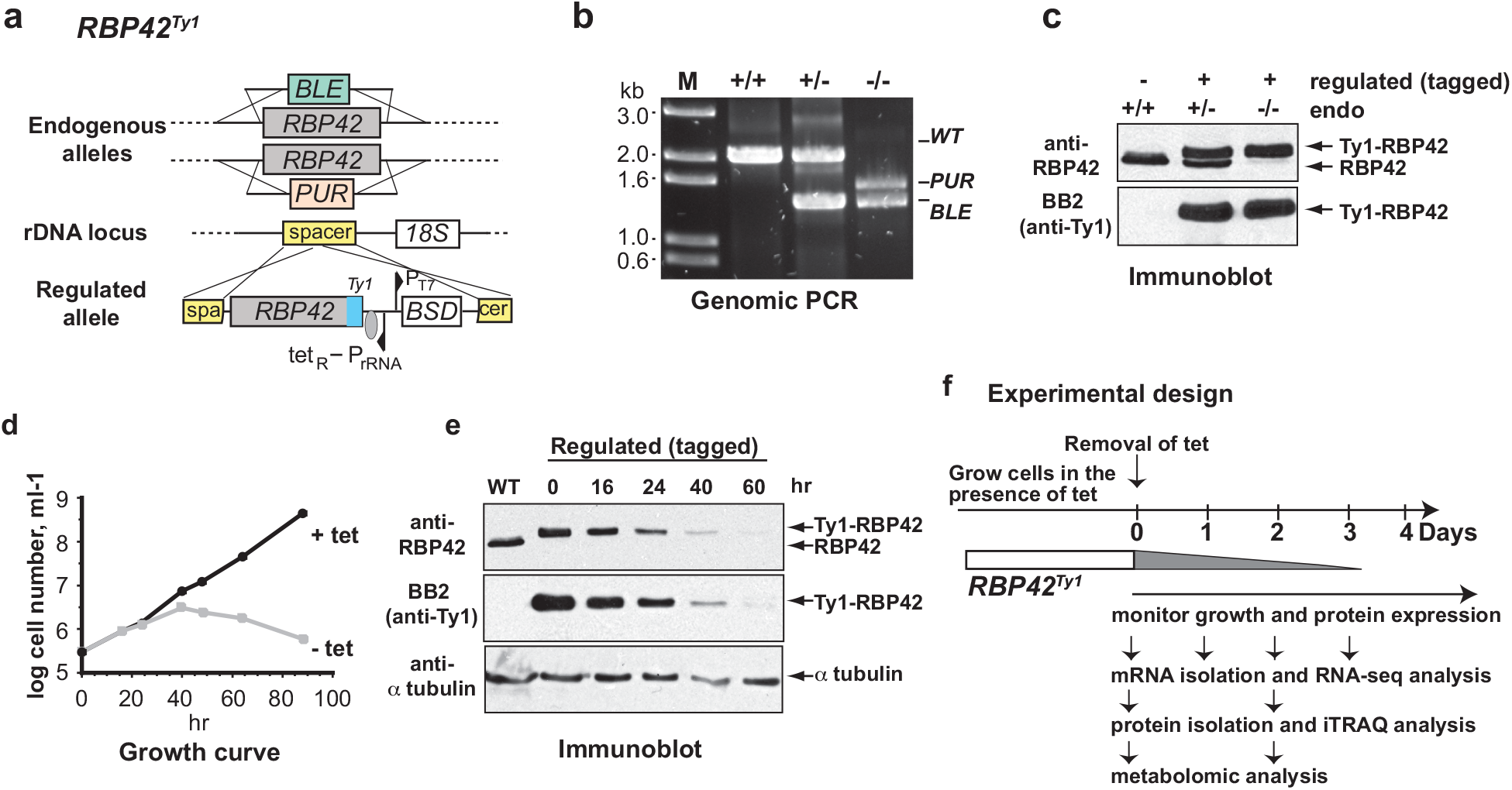
RBP42 is essential for slender bloodstream form *T. brucei*. **(a)** Schematic shows the strategy used to generate RBP42 conditional knockdown cell line (RBP42^Ty1^). Homologous recombination-mediated replacement by two antibiotic resistance genes inactivated two endogenous alleles. Cellular RBP42 expression is maintained from an exogenous tetracycline-inducible Ty1-tagged RBP42 allele inserted in an rRNA spacer region.**(b)** Analysis of genomic DNA by PCR, as in Figure 1, verifies proper integration of the antibiotic resistance DNA cassettes. **(c)** Immunoblot confirms expression of only the Ty1-tagged RBP42 protein from the regulated allele in RBP42^Ty1^ cells. Antibodies are described in Figure 1. **(d)** Growth analysis shows that RBP42 is essential for cell viability. RBP42^Ty1^ cells were maintained by adding tetracycline to the growth media. RBP42 expression was turned off by removing tetracycline from the growth media. Parasites were counted using a hemocytometer. **(e)** Immunoblot confirms regulated expression of the Ty1-tagged RBP42 protein in RBP42^Ty1^ cells. α-tubulin is loading control. **(f)** Schematic of experimental design. Expression of RBP42 was turned off on Day 0 by removing tetracycline from the growth medium. Knock down of RBP42 was indicated by gray ramp.

### Loss of RBP42 has minimal effect on target mRNA abundance

To determine the effect of RBP42 on its target mRNA stability, we measured transcriptome levels (mRNA-seq) in RBP42^Ty1^ cells before and after one, two, and three days of RBP42 depletion (Figure 3f). cDNA libraries, representing poly(A)^+^ RNA from triplicate samples of all four days were deep sequenced to obtain quantitative mRNA measurements (Figure S5a). Principal component analysis of mRNA-seq data shows robust clustering of samples from same day, and clear separations of clusters from all four days, confirming reproducibility of our measurement (Figure S5b). As expected, there is > 10-fold drop in RBP42 mRNA levels starting from day one (Figure S5c). Differential expression analysis, using DESeq2 ^38^, shows 208 mRNAs were significantly changed (adjusted p value < 0.01) following loss of RBP42: 111 upregulated (32 mRNAs > 2-fold), and 97 downregulated (only 2 mRNAs < 2-fold) (Supplementary Table). Upregulated mRNAs encode many cell-membrane associated proteins, including Variant Surface Glycoproteins (VSG) and membrane transport proteins; downregulated mRNAs encode a number of proteins involved in metabolic processes. However, loss of RBP42 did not elicit any clear, significant effect on abundances of its ‘most reliable’ mRNA target set (Figure S6a and b). Out of 183 mRNA targets 133 were slightly upregulated (to a maximum of 1.3-fold, 12 of them with adjusted p < 0.01), and 49 were marginally downregulated (to a minimum of 0.8-fold, 11 of them with adjusted p < 0.01) (Figure S6c). Therefore, we conclude that RBP42 is unlikely to control cytoplasmic turnover of its target mRNAs.

### Loss of RBP42 leads to decreased levels of target mRNA-encoded protein

To determine the effect of RBP42 on its target mRNA translation, we measured proteome levels in RBP42^Ty1^ cells before (day 0) and after two days of RBP42 depletion (day 2) using Isobaric Tags for Relative and Absolute Quantitation (iTRAQ) – based LC/MS/MS method. Total cellular proteins from eight samples, four replicates of each condition, were labeled with eight unique iTRAQ reagents and analyzed (Figure S7a). All eight samples exhibit similar iTRAQ intensity distribution, with a median value ∼100, indicating robust, reproducible measurement (Figure S7b). As expected, there is > 5-fold drop in RBP42 protein following two days of depletion (Figure S7c and d). Unbiased clustering of proteomes revealed that replicate samples of RBP42 depleted cells (day 2) cluster separately from replicate samples of normally growing RBP42 repleted cells (day 0), indicating reproducibility of iTRAQ quantitation (Figure S7e). Because of moderate to high positive correlation among replicate samples, analysis was performed combining replicate datasets. We observed major alterations in cellular proteome following loss of RBP42. Significant changes (p value < 0.01) were observed for 1650 proteins (∼30% of quantified proteome), of which 340 proteins showed ≥ 1.2-fold upregulation, and 226 proteins showed ≤ 0.8-fold downregulation (Figure 4a and Supplementary Table). Of the 183 ‘most reliable’ RBP42-target mRNA, identified by the iCLIP, our proteomic analysis quantified 178 mRNA-encoded proteins. Majority of the target-encoded proteins were downregulated; 109 proteins, of which 23 proteins are < 0.8-fold, in contrast to 69 upregulated proteins, of which 7 are > 1.2-fold (Figure S8). The noticeable changes in RBP42-target mRNA-encoded proteins following loss of RBP42, but not target mRNA abundance, indicate a possible translational regulatory role of RBP42 in *T. brucei* gene expression. Although it is possible that RBP42 exerts distinct translational regulations, both positive and negative, on discrete sets of mRNA targets, for the majority of its targets, RBP42 acts as a positive regulator of translation.

**Fig 4.**
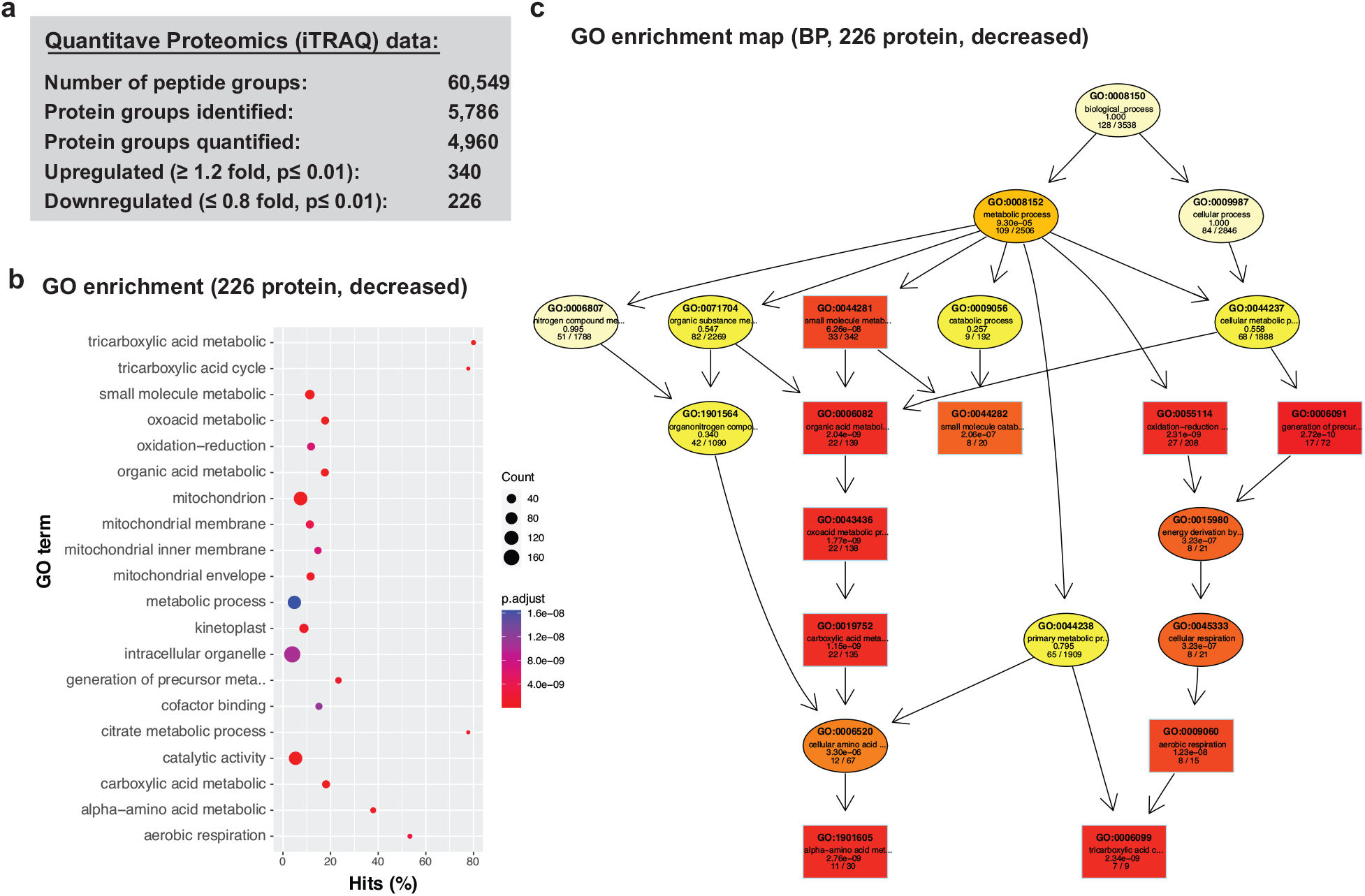
Loss of RBP42 reduces the levels of enzymatic proteins of central carbon and energy metabolic pathways. **(a)** iTRAQ quantitative proteomics of total cellular proteins from RBP42^Ty1^ cells before (Day 0) and two days after (Day 2) RBP42 knockdown. The number of protein groups with significantly (p ≤ 0.01, by paired sample t-test) changed levels (increased, ≥ 1.2-fold; decreased, ≤ 0.8-fold) are shown. **(b)** Gene Ontology (GO) enrichment analysis of 226 significantly down-regulated proteins following loss of RBP42. Dot plot showing top twenty enriched GO term. The x-axis represents percent hits, which is the ratio of the number of proteins in the 226 decreased set to the number of all annotated proteins with same GO term. The sizes of the dots represent the number of down-regulated proteins associated with the GO term. The colors of the dots represent adjusted p-values (BH). **(c)** GO graph showing significantly enriched GO terms in the Biological Processes category of the 226 down-regulated proteins (using TopGO). Each node marks a GO term, and each arrow indicates an ‘is-a’ relationship. Boxes indicate ten most significant nodes. Increasing coloring towards red represents increasing significance levels. The GO descriptions of each node along with significance levels and ratio of hits over total are also shown.

To evaluate the effects of loss of RBP42 on specific cellular processes, we performed GO term enrichment analysis of both up- and down-regulated proteins (Figure 4a). Many of the 340 most significant up-regulated proteins (> 1.2-fold, p < 0.01) are involved in membrane lipid and GPI anchor synthetic process, as well as membrane transporters, and show enrichment of a few general (higher level) GO terms. These include carbohydrate derivative biosynthetic process (GO:1901137, p 4.41e-7), and lipid metabolic process (GO:0006629, p 3.27e-6). In contrast, the 226 most significant down-regulated proteins (< 0.8-fold, p < 0.01) show many enriched GO terms, both general (higher level) and specific (lower level), with high significance (p < 1e-05) that are associated with primary metabolism (Figure 4b). Some very specific categories include tricarboxylic acid cycle (GO:0006099, p 2.34e-09), alpha amino acid metabolic process (GO:1901605, p 2.76e-09), and aerobic respiration (GO:0009060, p 1.23e-08). To obtain a meaningful biological interpretation, we analyzed these enriched GO terms using TopGO algorithm, which uses underlying GO graph topology to improve GO group scoring and reduce redundancy. The resulting GO graph of the Biological Process (BP) category, illustrated in Figure 4c, shows clear down-regulation of the central carbon and energy metabolic pathway following loss of RBP42.

To examine which specific processes are mostly affected, we compiled, using published dataset^23,25,39^, a set of gene cohorts associated with central carbon and energy metabolic pathway. Analysis shows down-regulation of many metabolic cohorts, but not control cohorts (Figure 5a). The two most affected cohorts are tricarboxylic acid cycle (TCA) and alpha amino acid oxidation. We observed significant down-regulation of enzymatic proteins that are part of the glycolytic, TCA cycle, and the alpha amino acid oxidation processes (Figure 5b). As anticipated, the levels of mRNAs encoding these enzymes did not show any significant changes (Figure S9). Although, existing metabolic model for slender bloodstream form does not support mitochondrial substrate-level or oxidative phosphorylation-mediated ATP production, recent proteomic studies revealed that most, if not all, enzymes involved in the production of succinate and acetate are expressed^40-42^. Importantly, several of these enzymes are also essential^40,43-44^. For example, we observed noticeable downregulation of these essential enzymes following loss of RBP42: PDH-E2 (Tb927.10.7570), 0.54-fold; TDH (Tb927.6.2790), 0.69-fold; α-KDE2 (Tb927.11.11680), 0.52-fold; and SCoAS β subunit (Tb927.10.7410), 0.55-fold. Identification of glucose-derived metabolic intermediates produced in the “succinate and acetate branches” of the TCA cycle in slender bloodstream form mitochondria further supports the activity of these enzymes ^41^. Downregulation of these essential pathways, therefore, is expected to perturb cellular central carbon metabolism.

**Fig 5.**
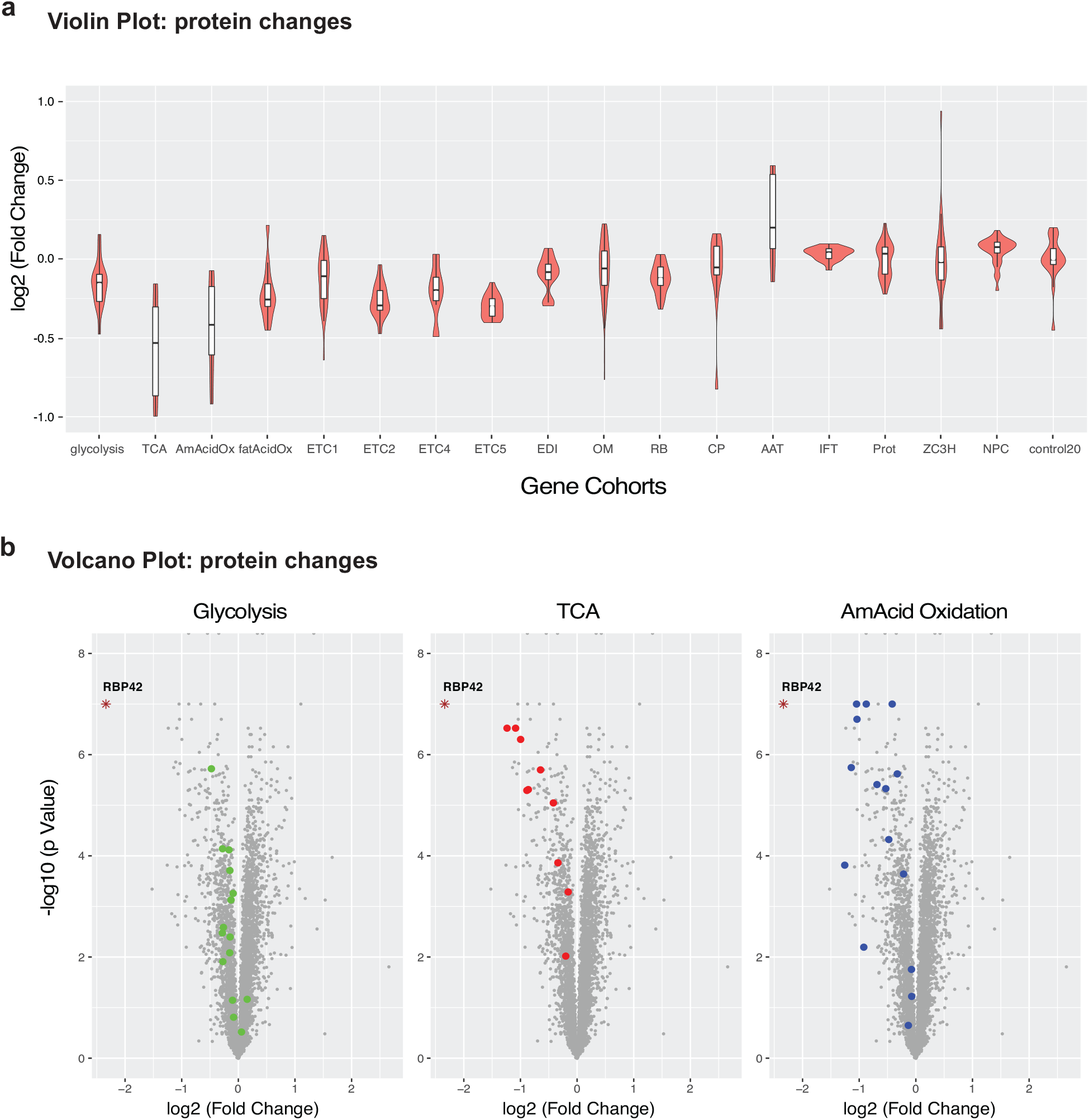
Loss of RBP42 causes significant downregulation of enzymatic proteins involved in glycolysis, TCA cycle, and amino acid oxidation. **(a)** Violin plots showing protein expression changes (iTRAQ intensity, log2) of gene cohorts following RBP42 knockdown. White box represents 25^th^ to 75^th^ percentile with the horizontal line as the median, and the whiskers extend 1.5 times the interquartile range. ETC1-5, mitochondrial respiratome complexes; EDI, editing complex; OM, mitochondrial outer membrane proteins; RB, RNA-binding complex; CP; carrier proteins; AAT, amino acid transporters; IFT, intraflagellar transport; Prot, proteasome; ZC3H, zinc finger family proteins; NPC, nuclear pore complex; control20, a cohort of 20 randomly sampled proteins. **(b)** Volcano plots showing changes in protein levels (log2 fold changes) versus significance p-values (-log10) for all (∼5000) quantified proteins (iTRAQ) after two days of RBP42 knockdown. Individual proteins that constitute glycolysis, TCA and alpha amino acid oxidation (AmAcid Oxidation) cohorts are color-coded.; all other proteins are shown in gray. Significance p values are estimated by paired sample t-tests.

### Loss of RBP42 impairs cellular central carbon and energy metabolism

To determine the effect of RBP42 on cellular metabolism, we measured levels of intracellular metabolites, including intermediates from glycolysis and TCA cycle, organic acids, and nucleotides. We reasoned that downregulation of the many metabolic enzymes following loss of RBP42 will alter the levels of these metabolic intermediates. Using mass spectrometry method, we generated metabolite profile of RBP42^Ty1^ cells before and after two days of RBP42 depletion. Compounds were extracted from four replicate samples of each condition and analyzed to measure quantitative changes in intermediary metabolites of glycolysis, TCA cycle, amino acids and nucleotides. Analysis revealed noticeable reduction in many glucose-derived metabolites; out of a total of 138 quantified metabolites, 40 show significant decrease (p < 0.05) (Figure 6a and Table S).

**Fig 6.**
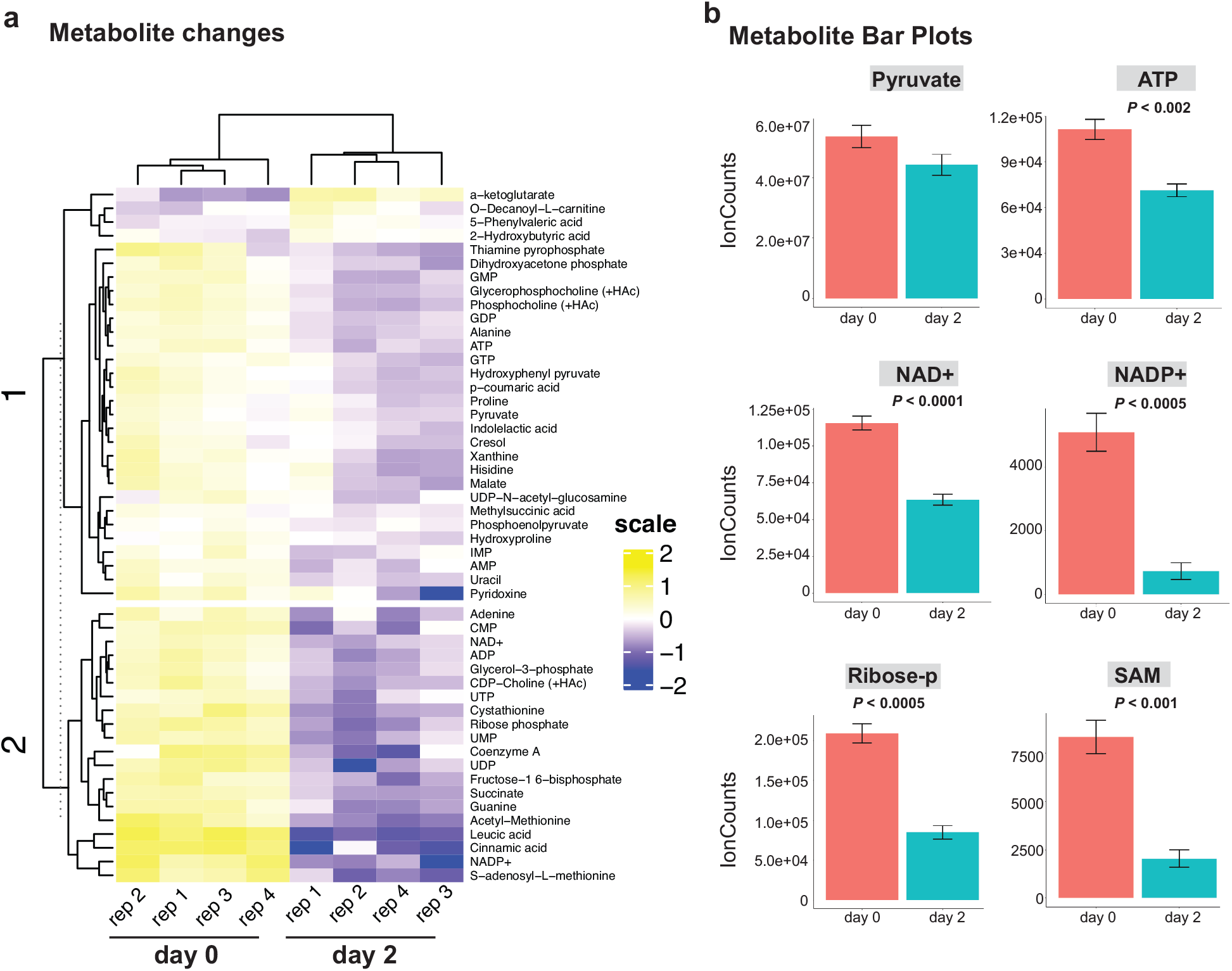
Loss of RBP42 alters cellular metabolic profile. **(a)** Heatmap view of metabolite levels following RBP42 knockdown. Metabolomes were quantified using LC-MS and metabolites were identified with known standards. Metabolites with significantly changed levels (p ≤ 0.05) are shown. Four replicate samples prepared from RBP42^Ty1^ cells before (day 0) and after two days of RBP42 knockdown (day 2) are indicated at the bottom. Metabolites are labeled on the right. The abundance of each metabolite is log2 transformed and mean-centered with blue being less abundant and yellow more abundant. The dendrograms are based on hierarchical clustering. **(b)** In the absence of RBP42 protein, several important molecules that determine cellular energy and redox states are significantly reduced. Bar plots showing mean ± standard error of signal intensities of six compounds before (red, day 0) and after (green, day 2) RBP42 knockdown. Significance p values, estimated by paired sample t-test are shown.

We observed noticeable (∼20%) decline in pyruvate, which is the major end product of catabolized glucose in slender bloodstream form. Pyruvate also serves as a major source of alanine, which declined 35% following loss of RBP42. We also observed large decreases in oxaloacetate, malate and succinate that are produced from the “succinate branch” of the TCA cycle. Oxaloacetate-derived aspartate is a known precursor of pyrimidine synthesis via dihydroorotate^45^. Significant decreases are also observed in purine and pyrimidine nucleotides. In the absence of de novo synthesis, trypanosomes rely on purine salvage^46^, which requires ribose 5-phosphate that is synthesized via the oxidative branch of pentose phosphate pathway^47^. We observed ∼60% decrease in ribose 5-phosphate, indicating that dysregulation of glucose metabolism also caused aberrations in pentose phosphate pathway. In addition to a steep decrease in ATP (∼40%), we observed significant decreases in critical cofactors, NAD and NADP, and the methyl donor S-adenosyl methionine, all of which are essential for normal cellular metabolic activities. Taken together, these results show that RBP42, by targeting mRNAs within the coding region, ensures proper regulation of core metabolic enzymes, which is critical for the parasite survival in the diverse nutritional environments encountered throughout its life cycle.

## Discussion

RNA-binding protein mediated post-transcriptional regulation play key roles in trypanosome gene expression. Although several important RNA-binding proteins have been studied to date, details of their regulatory roles remain elusive. Here, employing a detailed analysis that combines in vivo RNA target identification with global transcriptomic, proteomic, and metabolic profiling, we provide evidence that RBP42 acts as a critical regulator of *T. brucei* central carbon and energy metabolism in mammalian-infective slender bloodstream forms (Figure 7 and S10). Our analysis reveals that RBP42 targets mRNAs encoding enzymes involved in core metabolic processes. This finding is consistent with previously identified targets in the fly-infective procyclic forms^28^, indicating that RBP42 plays a conserved role in regulating metabolic genes in both stages of the parasite.

**Fig 7.**
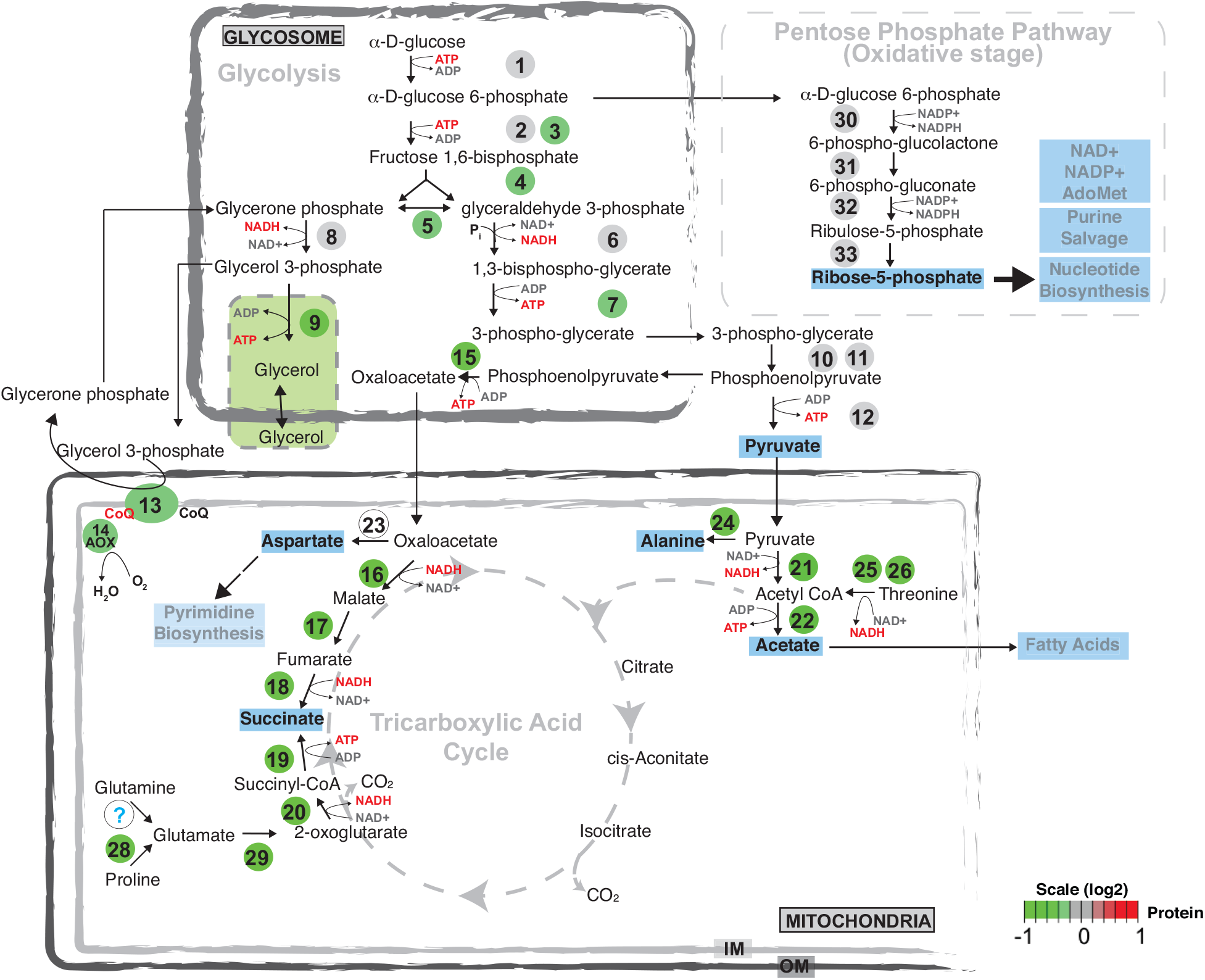
Changes in energy metabolic pathway following loss of RBP42. Schematic showing current consensus energy metabolic pathways that are active in slender bloodstream form trypanosomes. Dashed arrows indicate enzymatic steps for which no evidence of metabolic flux is available. Major intermediate metabolites that are either excreted out of the cell or utilized for biosynthesis of fatty acids or nucleotides are shown in bold. Numbered, colored circles indicate enzymes that are quantified in this study. Changes (log2 scale) in protein amounts are denoted using green-to-red color ramp. Enzymes are: 1, Hexokinase (HK); 2, Glucose 6-phosphate Isomerase (PGI); 3, Phosphofructo Kinase (PFK); 4, Fructose bisphosphate Aldolase (ALD); 5, Triose-phosphate Isomerase (TIM); 6, Glyceraldehyde-3-phosphate Dehydrogenase (GAPDH); 7, Phosphoglycerate Kinase (PGK); 8, NADPH-dependent glycerol-3-phosphate dehydrogenase (G3PDH, glycosomal); 9, Glycerol Kinase (GK); 10, Phosphoglycerate mutase (PGAM); 11, Enolase (ENO); 12, Pyruvate Kinase (PYK1); 13, FAD-dependent glycerol-3-phosphate dehydrogenase (G3PDH, mitochondrial); 14, Alternative Oxidase (AOX); 15, Phosphoenolpyruvate carboxykinase (PEPCK); 16, Malate dehydrogenase, mitochondrial (mMDH); 17, Fumarate hydratase, mitochondrial (FHm); 18, NADPH-dependent Fumarate reductase, mitochondrial (FRDm1); 19, Succinyl-CoA synthetase (SCoAS); 20, 2-Oxoglutarate dehydrogenase (2-OGDH); 21, Pyruvate dehydrogenase (PDH); 22, Acetate:Succinate CoA-transferase (ASCT); 23, Aspartate aminotransferase, mitochondrial (mASAT); 24, Alanine aminotransferase (ALAT); 25, Threonine dehydrogenase (TDH); 26, 2-Amino-3-ketobutyrate CoA ligase (AKCT); 28, Proline dehydrogenase, mitochondrial (PRODH); 29, Glutamate dehydrogenase (GDH); 30, Glucose-6-phosphate 1-dehydrogenase (G6PD); 31, 6-phosphogluconolactonase (6PGL); 32, 6-phosphogluconate dehydrogenase (6PGDH); 33, Ribose-5-phosphate isomerase (RPI).

RBP42 binds within the coding region of target mRNAs. Analysis of RBP42 crosslink sites on mRNA shows preference to a hexanucleotide sequence motif centered around a di-Cytidine (Figure 2d). However, a majority of the crosslink sites on RNA are devoid of this motif, raising the question of how RBP42 associates with specific sets of mRNAs. It is possible that RBP42 binds to cryptic RNA motifs, distributed on target mRNA after initial loading, or recruited to some target mRNA via protein-protein interactions.

Having defined RBP42’s RNA target set, we were motivated by curiosity to understand its impact on the regulation of these genes. We focused on the transcript and protein levels of the significant subset of iCLIP-identified targets following loss of RBP42, using a conditional knockdown strategy. Depletion of RBP42 had little effect on steady state levels of target mRNAs, but major effect on the target proteome, indicating a possible translational regulatory role of RBP42. This is consistent with polysomal association of RBP42 observed in a previous study^28^.

Our analysis indicates that loss of RBP42 caused clear downregulation of many enzymes involved in core metabolic processes. Along with glycolysis, the two most affected pathways are mitochondria-resident interconnected energy metabolic pathways, i.e., TCA cycle and amino acid oxidation. It is increasingly recognized that many cellular mRNAs are translated within subcellular domains that allows precise localization and regulation of the newly made proteins^48^. RBP42 may permit translocation of mitochondrial-resident proteins synthesis, while protecting mRNA from degradation. Similar coordinated control of mitochondrial respiratome components is proposed for *T. brucei* ZC3H39/40 RNA-binding complex^25^. RBP42 is known to be phosphorylated^49^, which is a well-recognized mechanism that control activity of many RBPs^50-52^. It is possible that phosphorylation of RBP42 is regulated by nutrient availability, which in turn may regulate its activity as a translational regulator to fine tune cellular metabolic activity.

Importantly, loss of RBP42 resulted in significant reduction of many intermediary metabolites. Energy metabolism, i.e., production and utilization ATP, is a highly complex, dynamic process involving numerous factors that respond to intra- and extra-cellular signals, nutrient availability, and cellular physiological and developmental status. Slender bloodstream form *T. brucei* relies mainly on glucose as the major carbon source and current metabolic model suggests that ATP is exclusively produced via glycolysis, pyruvate being the major (∼85%) end product^41^. The high ATP demand of proliferating cells is achieved by ∼10-fold greater glycolytic rate in slender bloodstream form, compared to procyclic form. However, recent proteomic studies have confirmed that although mitochondrial oxidative- and substrate level-phosphorylation is not utilized for ATP production, various mitochondrial activities are essential for slender form survival. Metabolomic studies confirmed that slender form excretes significant levels of alanine, acetate and succinate that are produced in the mitochondria. Mitochondrial production of acetate is essential for the slender bloodstream form *T. brucei*^40^. Glucose derived pyruvate and threonine are two main sources of acetate, which is produced via the “acetate branch” of TCA cycle. Therefore, marked reduction of both pyruvate dehydrogenase (PDH) and threonine dehydrogenase (TDH), following RBP42 knockdown, is expected to severely weaken acetate production.

RBP42 knockdown reduced the levels of many glycolytic enzymes. This modulation of glycolysis may impede proper function of the oxidative branch of pentose phosphate pathway and therefore reduce ribose 5-phosphate and nucleotide salvage pathways. A marked, ∼ 40%, drop in ATP level was observed. Cellular metabolic activities primarily rely on ATP as energy carrier that drives anabolic reactions critical for cell survival and proliferation. Importantly, ATP also works as structural precursor of important cellular cofactors, including NAD, NADP and the methyl donor S-adenosyl methionine (SAM). Therefore, a decrease in ATP level is expected to cause widespread disruption of metabolic activity. For example, a decline in SAM is expected to cause major alterations in activities of many methylation-dependent RNA-binding protein. Absence of protein arginine methylation is known to cause striking changes in cellular energy metabolism^27^.

In conclusion, it is clear that the RBP42 allows proper expression of metabolic enzymes involved in *T. brucei* central carbon and energy metabolism. By undertaking a broad approach, we show that RBP42 mediated regulation of metabolic networks is critical for the parasite. Although RBP42’s precise mode of action remains to be discovered, our analysis hints an apparent translational regulatory role of RBP42.

## Materials and Methods

### *T. brucei* strain and growth analysis

*T. brucei* Lister 427 bloodstream form wild-type cells, single marker (SM) cells^53^, and all stable transgenic cell lines were grown in HMI-9 medium supplemented with 10% fetal bovine serum and 10% serum plus at 37°C in a humidified incubator containing 5% CO2. SM cells that co-express T7 RNA polymerase and the Tet repressor with *NEO* resistance gene are maintained with 2.5 μg/ml G418. Recombinant DNA constructs were introduced by nucleofection using Amaxa Human T-solution following manufacturer’s instruction. Transgenic cell lines were selected and grown by the addition of puromycin, phleomycin and blasticidin as required at 0.1, 1.25 and 5 μg/ml, respectively. Homozygous triple-Ty1-tagged RBP42 cell line was generated by N-terminal epitope tagging of both RBP42 alleles at the native loci. RBP42 conditional knockdown cell line (RBP42^Ty1^) was generated by replacing two native RBP42 alleles with *BLE* and *PAC* selectable marker cassettes and introducing an ectopic inducible Ty1-tagged RBP42 allele. The inducible expression construct was introduced into a single RBP42 allele null strain prior to deletion of the second RBP42 allele, while maintaining RBP42 expression by addition of tetracycline to the culture media. RBP42^Ty1^ cells were grown in the presence of 1 μg/ml tetracycline to maintain inducible expression of exogenous RBP42 transgene. To shut down expression of the exogenous RBP42 transgene, cells were washed twice to remove tetracycline and resuspended in culture medium lacking tetracycline. Cultures were seeded at 1 × 10^5^ cells/ml and growth analysis was carried by counting cell density on a hemocytometer.

### Plasmid constructs

Epitope tagging of RBP42 alleles at the native loci was carried out using PCR-based strategy^54^. Two PCR-generated DNA modules, one with *NEO* resistance gene and the other with *PAC* resistance gene, were used in two successive rounds of transfection and selection to tag two native RBP42 alleles. The ectopic inducible construct was made by inserting Ty1-tagged RBP42 open reading frame (ORF) into plasmid pAD74 ^55^. The resulting construct was linearized using NotI restriction enzyme to facilitate homologous recombination into an rRNA loci. A *BLA* resistance gene within the construct allowed selection of stable cell lines. To replace endogenous RBP42 alleles, knockout gene cassettes were generated by cloning 500 bp upstream and 1000 bp downstream sequences to flank *BLE* and *PAC* resistance genes. Knockout cassettes were released by restriction enzyme digestion prior to transfection.

### Crosslink immunoprecipitation (CLIP)

CLIP was carried out with minor modifications of previously published method^28^. Briefly, slender bloodstream forms (0.5 to 0.8 × 10^6^ cells/ml) were harvested and washed once with cold trypanosome buffer saline (TBS: 137 mM NaCl, 2.7 mM KCl, 10 mM Na2HPO4, 1.8 mM KH_2_PO_4_, 20 mM glucose, pH 7.4). Cells, resuspended in 6 ml TBS to a concentration of ∼5 × 10^7^ cells/ml, were transferred to a 100-mm Petri dish, placed on an ice tray, and UV-irradiated (254nm) once with 150 mJ/cm^2^ or 300 mJ/cm^2^ dose in Stratalinker 1800 (Stratagene) UV light source. Cells were rapidly pelleted, snap frozen in liquid N2, and stored in -80°C in small aliquots. Crosslinked RBP42-RNA complexes were immunopurified from cellular extracts using antibodies attached to magnetic beads (Dynabeads): anti-RBP42 antibody to Protein G beads and anti-Ty1 (BB2) antibody to sheep anti-mouse IgG beads. Captured protein-RNA complexes were washed extensively and analyzed by gel chromatography.

### iCLIP-Seq

iCLIP-Seq libraries were prepared following published method^56^. Prior to immunoprecipitation, cellular extracts were treated with titrated amount of RNase I to generate median 40-80 nucleotides (nt) long RNAs. After high-salt stringent washing, RNAs on beads were ligated to an adapter at the 3’ end, and radioactively labeled on the 5’ end. Protein-RNA complexes, run on 4-12% NuPAGE™ Bis-Tris gel (Invitrogen) and transferred on to Nitrocellulose membrane, were detected using X-ray film. Complexes in the range of 20-40 kDa above RBP42 were excised, and RNA was recovered by proteinase K digestion. Reverse transcription of RNA was performed using primers with two cleavable adapter regions separated by a *Bam*HI site, as well as a barcode to mark unique cDNA molecules. cDNAs were size-selected into two sizes, 80-100 nt as low (L) and 100-150 nt as high (H), using denaturing gel electrophoresis, circularized by CircLigase™ II ssDNA ligase (Lucigen). Subsequently, cDNAs were linearized by BamHI digestion, PCR-amplified, and sequenced on the Illumina NextSeq platform.

iCLIP-Seq data were analyzed using published ‘pipeline’^57^. Raw sequence reads, quality checked using FastQC (www.bioinformatics.babraham.ac.uk/projects/fastqc), were trimmed to remove adapter and barcode sequences. The barcode sequence was assigned to each read, using Flexbar, as unique molecular identifier (UMI) that was later used to eliminate PCR duplicates^58-59^. Sequence reads of at least 15 nt in length were mapped to *T. brucei* 927 genome assembly version 45 using STAR aligner with parameters set to search only unique alignment, with less than 4% mismatched bases^60^. Following removal of PCR duplicates using UMI-tools, each unique cDNA molecule was counted as an independent crosslink event^59^. Crosslinked sites, the nucleotide preceding the cDNA start, were extracted using BEDTools suit^61^. Significant crosslink sites were determined using PureCLIP cluster finding algorithm^37^. Crosslink sequences were extracted by adding 5 nt flanking sequences to crosslink sites to obtain 11 nt crosslinking regions. Crosslink sequence motif discovery was performed using the MEME suit (meme-suite.org/tools/meme), with the parameters -classic mode, zero or one occurrence per sequence, search given strand only, minimum width 4, maximum width 10.

### mRNA-Seq

mRNA-Seq libraries were prepared from poly(A)^+^-containing RNA, captured by two rounds of oligo-d(T)n-bead selection of 10 μg total RNA, following Illumina small RNA library preparation method. Libraries were sequenced on the Illumina NextSeq platform. Sequence reads were mapped to the *T. brucei* 927 genome assembly version 45 using Bowtie2 (v2.3.5). The mapped reads were then converted to gene expression values and analyzed using DESeq238. Volcano and violin plots were generated using expression value (log2-fold change) and p-value in R.

### iTRAQ quantitative proteomics

Quantitative proteomics using iTRAQ method were performed in the Center for Advanced Proteomics Research (NJMS, Rutgers Biomedical and Health Sciences)^62^. RBP42^Ty1^ cells (5 × 10^7^ cells/assay) were harvested before (day 0) and after two days (day 2) of RBP42 knockdown, and washed with ice-cold TBS. Total proteins were extracted using lysis buffer containing 100mM TEAB, 8M urea and protease inhibitor cocktail. One hundred microgram proteins, from four replicate samples of each condition, were reduced, alkylated, and trypsin digested before subjected to labelling with 8-plex iTRAQ reagents (AB Sciex). Peptides from day 0 replicates were labeled with iTRAQ tag-113, 114, 115 and 116, whereas peptides from day 2 replicates were labeled with iTRAQ tag-117, 118, 119 and 121. Subsequently, all labeled peptides from 8 samples were pooled and fractionated using high pH RPLC liquid chromatography on ACQUITY UPLC system (Waters Corporation). A total of 48 fractions were collected in 60-min gradient of Solvent A (20 mM HCOONH_4_, pH10.0) and Solvent B (20 mM HCOONH_4_ in 85% ACN, pH10.0) and pooled into 12 fractions that were subjected to LC-MS/MS analysis on an UltiMate 3000 RSLCnano coupled with Orbitrap Fusion Lumos Mass Spectrometer (Thermo Scientific). Peptides, ∼1 µg from each fraction, were separated on a nano C18 column (Acclaim PepMap, 75µm × 50cm, 2µm, 100Å) using a 2-hour non-linear binary gradient of mobile phase A (2% ACN and 0.1% formic acid) and mobile phase B (85% ACN and 0.1% formic acid) at a flow rate of 300 nl/min. Eluted peptides were introduced into Orbitrap Fusion Lumos system through a nanospray Flex™ ion source (Thermo Scientific) with the spray voltage of 2kV and a capillary temperature of 275°C. The MS spectra was acquired in a positive mode. For MS1, peptide scan range was set to 375-1,500 with the resolution of 120,000. Peptides with charge-state of 2-7, and intensity greater than 5 × 10^3^ were selected for MS/MS scan in ion-trap using collision-induced dissociation (CID) with the collision energy of 35%. The dynamic exclusion is 60s and the isolation window is 0.7 m/z. For SPS-MS3 scan, the precursor selection range was 400-1200 with iTRAQ ion excluded. Ten SPS precursors were selected for MS3 scan in orbitrap with resolution 50,000. High energy collision dissociation (HCD), with the collision energy of 65%, was used for iTRAQ tag quantitation.

The iTRAQ MS data were searched against UniProt *Trypanosoma brucei brucei* (strain 927/4 GUTat10.1) database (8579 proteins) using Sequest search engine on Proteome Discoverer (V2.4) platform. MS1 mass tolerance was set to 10ppm and MS2 mass tolerance was 0.6Da. iTRAQ 8-plex (K), iTRAQ 8-plex (N-terminal) and methylthio (C) were set as fixed modification, whereas oxidation (M) and iTRAQ 8-plex (Y) as variable modifications. Two missed cleavages are allowed in trypsin digestion. The reporter ion-based quantification workflow was chosen for data analysis; the CID spectra in MS2 was used for peptide identification and the HCD spectra in MS3 was used for iTRAQ quantitation. The false discovery rate for protein and peptide were set to 1% filtered with Percolator. The protein relative quantitation was calculated based on the ratio of (average in day 2 abundance)/(average in day 0 abundance). Significance (p values) is computed using paired sample t-test.

### Metabolomics

Metabolomic profiling was performed in the Metabolomics Shared Resources facility (Rutgers Cancer Institute of New Jersey), following published method^63^. Briefly, cells (2 × 10^7^ cells/assay) were harvested as above. Metabolites were extracted with 1 ml of 40:40:20 mixture of methanol:acetonitrile:water plus 0.5% (V/V) formic acid on ice for 5 min. Following neutralization of formic acid by addition of 50 μl of 15% (m/V) NH4HCO3, cleared extracts were collected by centrifugation at 15000g for 10 min and stored at -80°C. Metabolomic data, designed to capture intermediary metabolites in central carbon metabolism including glycolytic intermediates, TCA compounds, amino acids, nucleotides and derivatives, were obtained using hydrophilic interaction liquid chromatography (HILIC) separation method coupled with mass spectrometry run in negative ionization mode. Each metabolite was identified by matching of accurate mass and retention time to synthetic standards. Metabolite bar plots were generated using mean ion counts ± standard errors. Significance (p value) is computed using paired sample t-test.

### Flow cytometry

Cells (1 × 10^7^ cells/assay) were stained with Hoechst 33342 (1 μg/ml) in growth medium at 37°C for 10 min and run on a BD-LSR (Becton Dickinson, NJ USA). Data was analyzed using FlowJo.

### Fluorescence Microscopy

Cells were fixed in 1% formaldehyde and adhered to slides coated with poly L-Lysine. RBP42 immunolocalization was performed using anti-RBP42 antibodies at 1:10,000 on cells permeabilized with 0.2% NP40 for 5 min at room temperature. FITC conjugated secondary antibodies are used at 1:1000. Cells were mounted with Vectashield (Vector Laboratories) containing DAPI and imaged using an Olympus BX61 microscope equipped with DAPI and FITC-sensitive filters and a Hamamatsu ORCA-ER camera.

### Gene Ontology (GO) and pathway enrichment analysis

GO term enrichment analysis was performed using resources available from TriTrypDB.org database. REVIGO web tool was employed to summarize GO enrichment by eliminating redundant GO terms^64^. Directed acyclic graph of enriched GO terms was produced using R Bioconductor topGO package^65^. Pathway analysis was performed by R Bioconductor packages GAGE^66^ and Pathview^67^ using Kyoto Encyclopedia of Genes and Genomes (KEGG) database^68^.

