## Supplemental Figures for "The RNA-binding protein RBP42 regulates cellular energy metabolism in mammalian-infective *Trypanosoma brucei*"

a

iCLIP seq data

| cDNA numbers | Sample 1 | Sample 2 | Sample 3 | Sample 4 |
|---|---|---|---|---|
| Raw reads | 29,576,322 | 11,326,474 | 15,397,518 | 35,773,106 |
| Mapped (STAR) | 26,600,252 | 9,842,221 | 12,694,064 | 29,501,373 |
| % Mapped | 89.93% | 86.89% | 82.44% | 82.46% |
| Uniquely mapped | 16,594,863 | 6,744,936 | 6,965,986 | 15,918,272 |
| % Uniquely mapped | 56.11% | 59.55% | 45.24% | 44.50% |
| Unique cDNA molecule | 29,464 | 26,055 | 867,214 | 493,361 |
| Reads with deletion | 513 | 569 | 40,436 | 23,881 |
| Mapped to genes | 26,853 | 21,811 | 814,291 | 453,422 |
| Mapped to CDS | 22, 483 | 15,877 | 723,370 | 425,516 |
| Mapped to intergenic (5' and 3' UTR) | 4,175 | 5,839 | 72,987 | 28,731 |
| Mapped to tRNA, rRNA, other snRNA | 123 | 88 | 15,318 | 7,113 |
| Mapped to No feature | 2,533 | 4,122 | 51,043 | 39,173 |

b

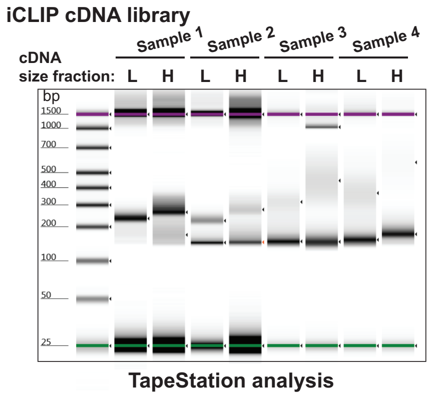


c

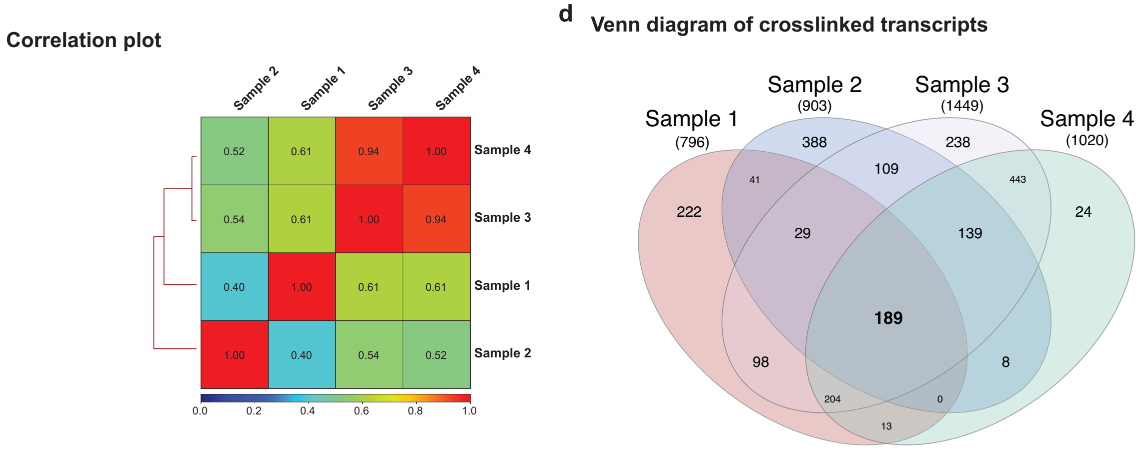


**Fig S1. Details of high-throughput sequencing of iCLIP cDNA libraries**

**(a)** Illumina iCLIP-seq data showing cDNA tag counts. Numbers of unique cDNA molecules identified in each sample are highlighted in yellow. **(b)** Agilent 2200 TapeStation system Screen Tape gel image of PCR amplified cDNA libraries. Prior to PCR amplification, CLIP purified cDNAs from each sample were size fractionated into two, H, high and L, low, by gel chromatography. Note that in samples 3 and 4 (with 300mj/cm$^2$ UV dose) amplified cDNAs are of varied length and form diffused bands compared to compact bands visible in samples 1 and 2 (with 150mJ/cm$^2$ UV dose). High-throughput sequencing was performed by pooling both H and L fractions from each sample. **(c)** All four iCLIP samples shows moderate to high correlations. Clustered heatmap of pairwise Spearman correlations of unique cDNA reads are shown. cDNA reads, covering every 10,000-base regions in the genome, are counted, normalized, and compared (using deeptools). **(d)** Venn diagram of target transcripts identified in all four iCLIP cDNA libraries. The 189 congruent targets present in all four samples are in bold.

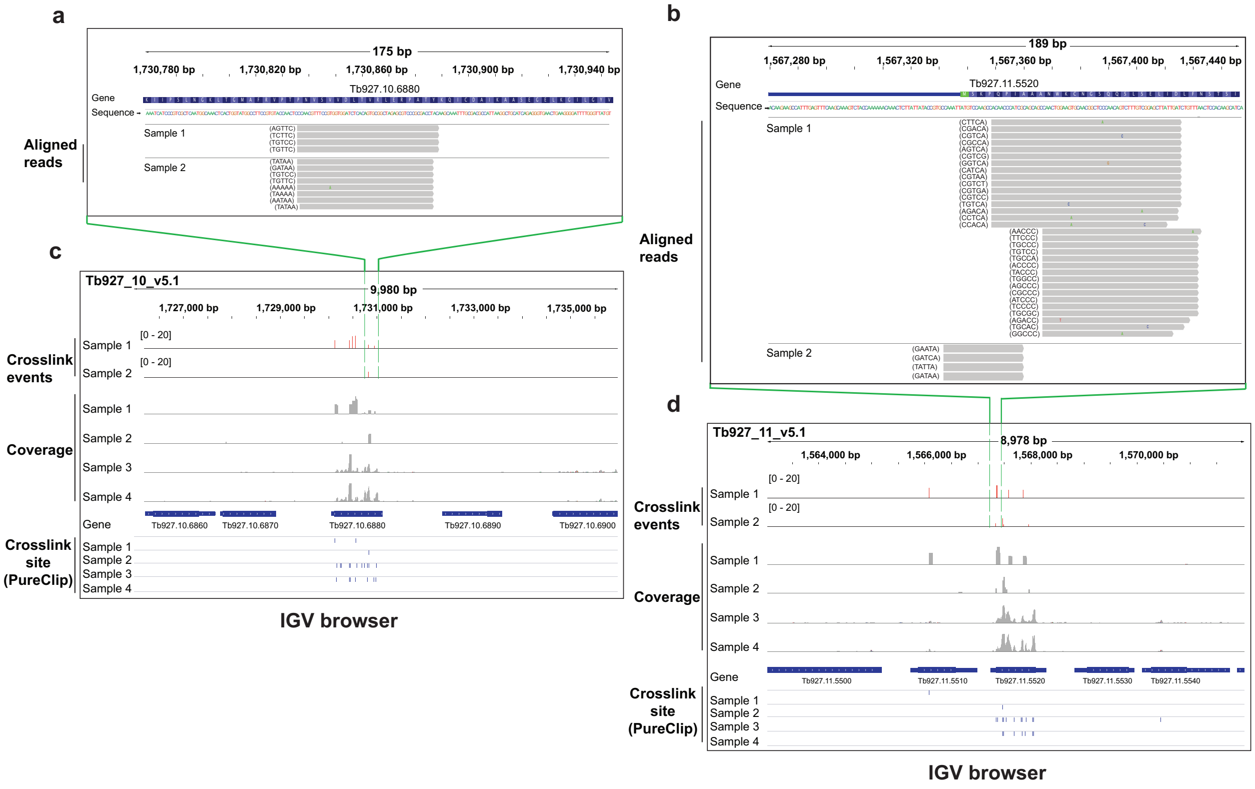


**Fig S2. IGV browser view of mapped cDNA read clusters on representative crosslink sites**

**(a) and (b)** The genomic and the protein amino acid sequences of two ~ 180 bp chromosomal loci that are part of two target genes (Tb927.11.5520 and Tb927.10.6880) are shown. The 5nt unique molecular identifier (UMI) sequences of stacked coinciding cDNAs are indicated in parenthesis. **(c) and (d)** Details of iCLIP data, presented in Figure 2c, showing two ~ 9 kb genomic loci on chromosomes 11 and 10. In the top panel, the red bars indicate the number of crosslink events on crosslink sites (see Figure 2c). In the middle panel, gray histograms show estimated read coverages (normalized, reads per million) for all four samples. In the bottom panel, the blue ticks mark the crosslinking sites predicted by PureCLIP algorithm.

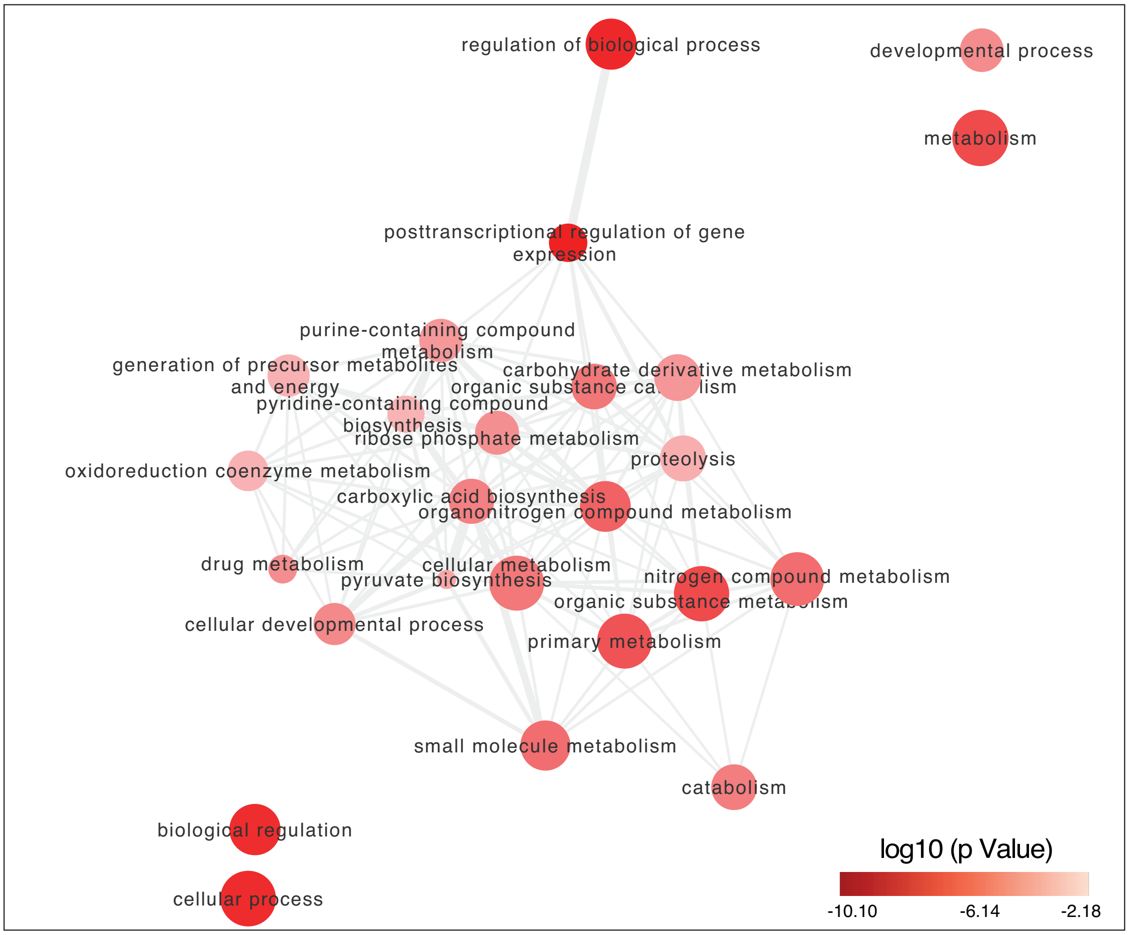


**Fig S3. Gene Ontology (GO) terms enrichment analysis**

A graph view of enriched Gene Ontology (GO) terms, associated with the 165 annotated RBP42 target genes, generated using REVIGO webtools (http:// revigo.irb.hr/) and visualized using Cytoscape program. The bubble radius indicates generality of the GO term. The color indicates significant enrichment p-values

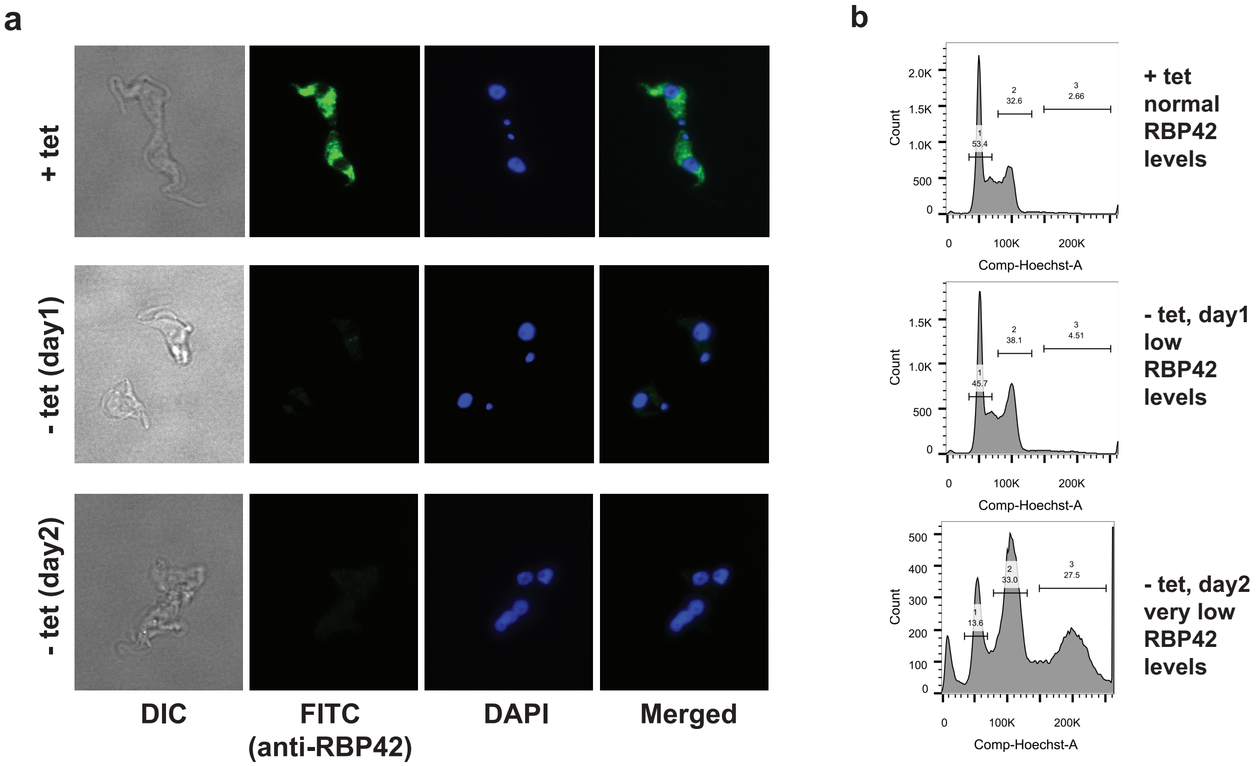


**Fig S4. Phenotypic effect of RBP42 knockdown on slender bloodstream form *T. brucei*** **(a)** Immunofluorescence microscopy shows robust expression of RBP42 protein, as expected, when RBP42$^{Ty1}$ cells are grown in the presence of tetracycline (+ tet, top panel). However, RBP42 protein is not detectable after removal of tetracycline (- tet, day1 and day2, middle and bottom panel). Also visible are multinucleated cells (- tet, day2, bottom panel), similar to the phenotype observed in procyclic *T. brucei* (Das et al. 2012). **(b)** Flow cytometry analyses show that as parasites lose RBP42, cells become blocked in G2 stage of the cell-cycle. 1 is 1n, cells with normal amount of DNA, value of '50k' for Hoechst staining; 2 is 2n, value of '100k' (50 x 2); 3 is > 2n with a peak value of '200k' (50 x 4). Percentage of cells under the peak regions are shown. Parasites in the region 3 consist of more than two nuclei, which indicates loss of proper cytokinesis.

**a** mRNA seq data

| Samples | Replicates | Number of reads | % mapped | % uniquely mapped |
|---|---|---|---|---|
| Day0 | Rep1 | 60,230,449 | 93.05 | 65.86 |
| | Rep2 | 57,881,419 | 93.14 | 66.10 |
| | Rep3 | 52,117,872 | 93.19 | 65.46 |
| Day1 | Rep1 | 54,468,139 | 92.53 | 63.97 |
| | Rep2 | 44,753,512 | 92.66 | 62.77 |
| | Rep3 | 52,975,957 | 92.63 | 63.89 |
| Day2 | Rep1 | 45,176,389 | 93.17 | 66.42 |
| | Rep2 | 40,440,496 | 93.18 | 64.77 |
| | Rep3 | 41,980,664 | 92.99 | 64.42 |
| Day3 | Rep1 | 34,083,506 | 93.20 | 65.54 |
| | Rep2 | 37,599,589 | 93.32 | 66.26 |
| | Rep3 | 33,299,932 | 93.34 | 66.56 |

**b** PCA plot

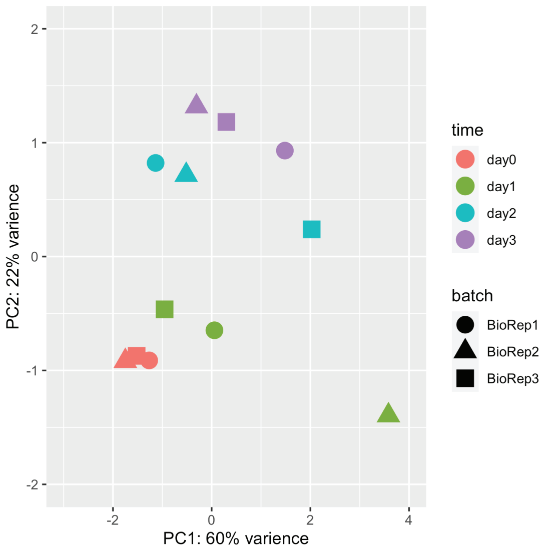


**c** mRNA levels

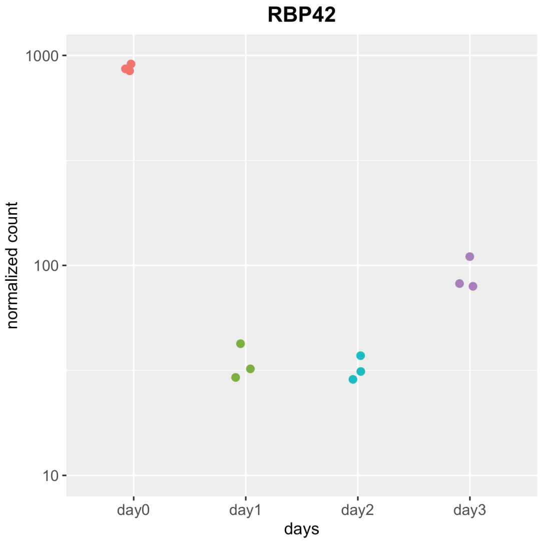


**Fig S5. Effect of RBP42 knockdown on global transcriptomes (mRNA levels) of slender bloodstream form *T. brucei***

**(a)** Illumina mRNA-seq data from RBP42$^{Ty1}$ cells before (Day 0) and after (Days 1-3) RBP42 knockdown. The table shows total number and percent of uniquely mapped reads from three replicate samples of each. **(b)** Principle component analysis (PCA), performed with rlog-normalized expression data (DESeq2), reveals separation of clustered samples belonging to different 'day'. Replicate samples from each day are color coded. **(c)** Dot plot of normalized count (DESeq2) showing efficient (> 10-fold) reduction of RBP42 mRNA. Estimated RBP42 mRNA levels of all twelve samples are shown. Color code as in (b).

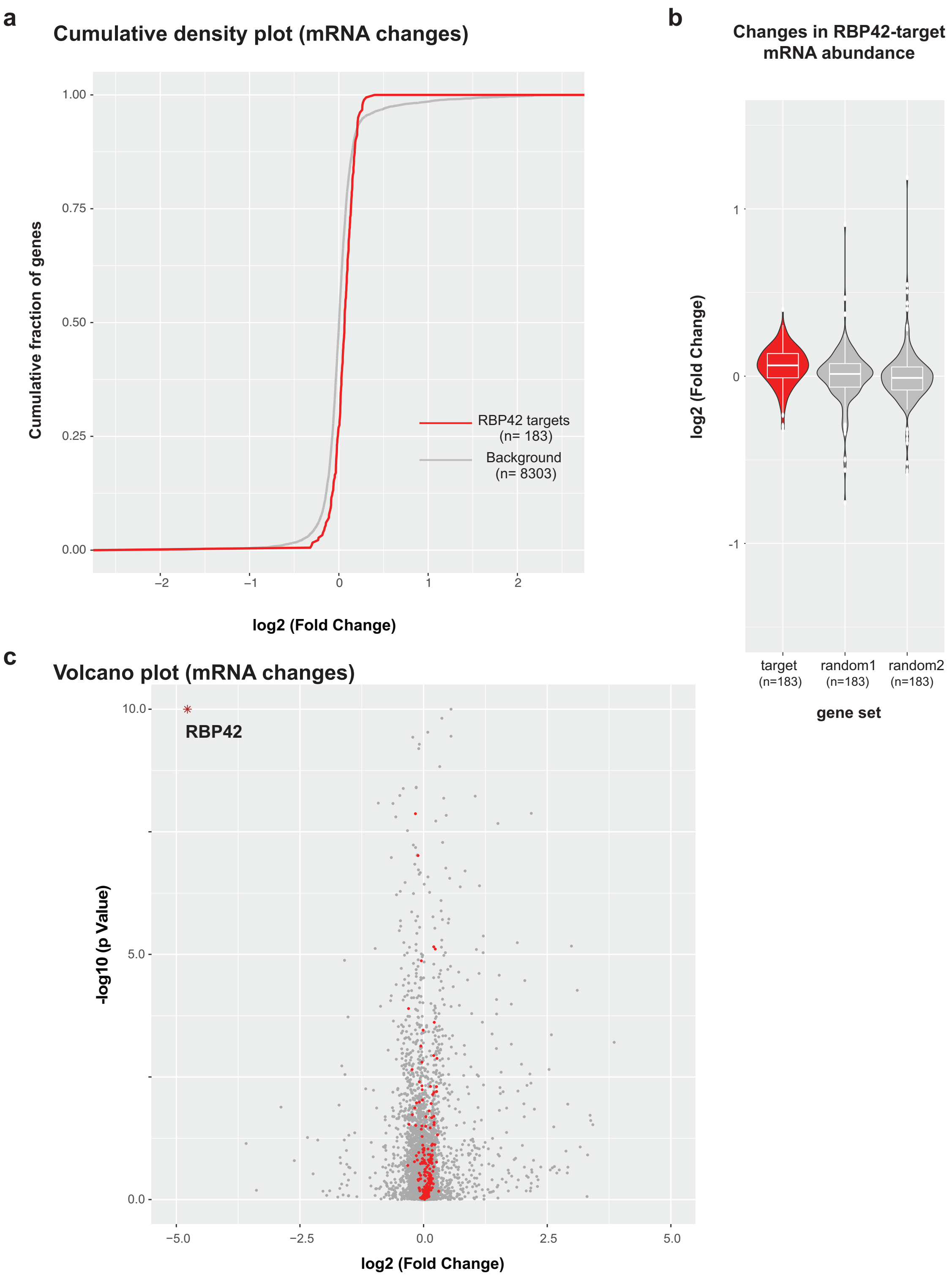
a
Cumulative density plot (mRNA changes)
Cumulative fraction of genes
1.00
0.75
0.50
0.25
0.00
−2
−1
0
1
2
log2 (Fold Change)
RBP42 targets
(n= 183)
Background
(n= 8303)
b
Changes in RBP42-target
mRNA abundance
log2 (Fold Change)
1
0
-1
target
(n=183)
random1
(n=183)
random2
(n=183)
gene set
c
Volcano plot (mRNA changes)
-log10 (p Value)
10.0
5.0
0.0
RBP42
−5.0
−2.5
0.0
2.5
5.0
log2 (Fold Change)

**Fig S6. Loss of RBP42 has minimal effect on its target mRNA abundance**

**(a)** Cumulative distribution frequency (CDF) plot shows the empirical cumulative distribution of log2 fold changes for RBP42-targets (red) compared to non-target background (grey) following RBP42 knockdown. The almost overlapping CDF of the RBP42-targets (red) with the background (grey) indicates no major changes in mRNA levels. **(b)** Violin plots showing the distribution of fold changes (log2) of the 183 target mRNAs (target, red). Similar plots of two equal-numbered control gene sets, randomly sampled (random1 and random2, grey) from the mRNA-seq data, are also shown for comparison. White box represents $25^{th}$ to $75^{th}$ percentile with the horizontal line as the median, and the whiskers extend 1.5 times the interquartile range. **(c)** Volcano plot showing changes in mRNA levels (log2 fold changes) versus p-values (-log10) for all (~8500) protein coding genes (mRNA-seq) after two days of RBP42 depletion. The set of 183 RBP42-targets are highlighted in red; all other genes are shown in grey.

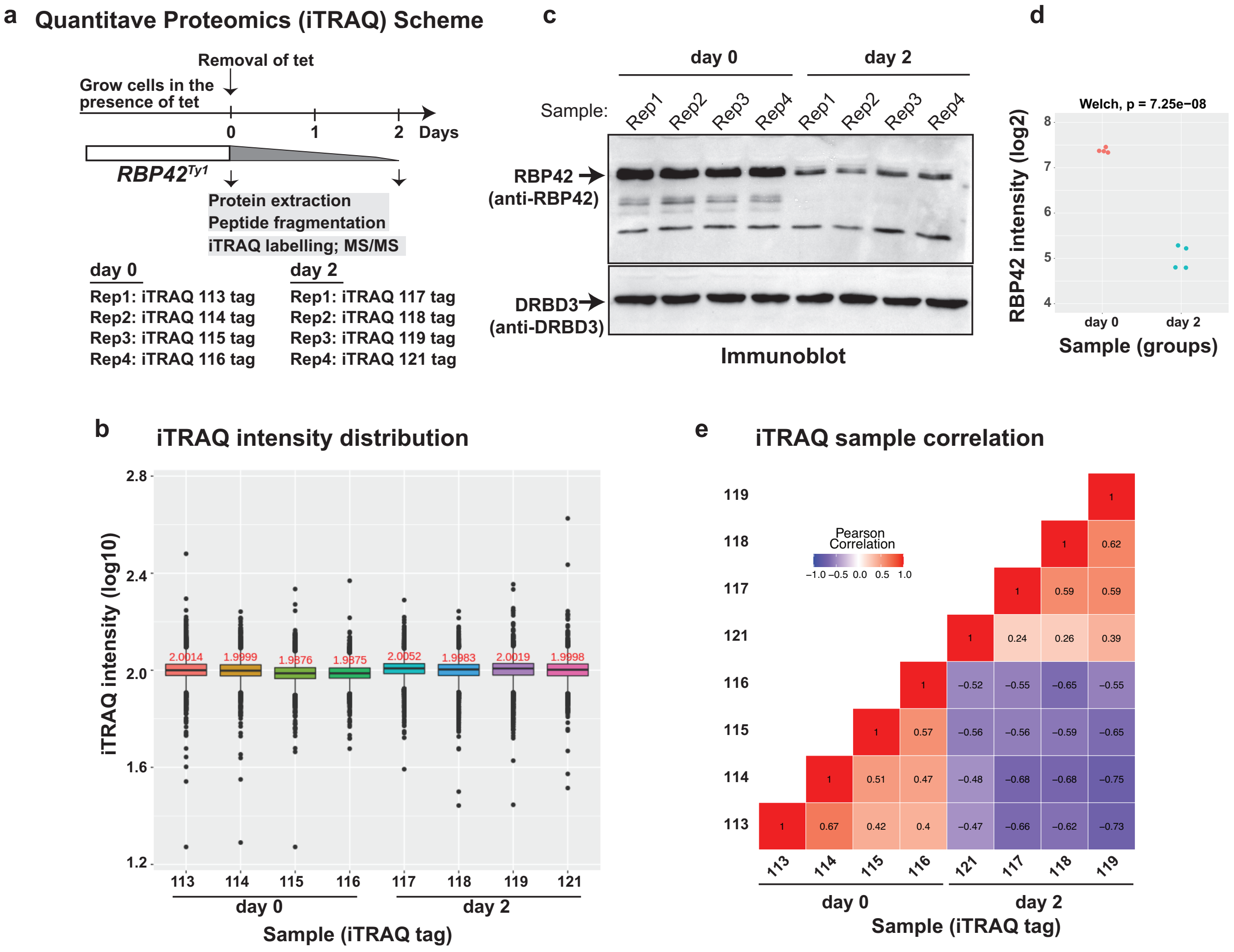


**Fig S7. Effect of loss of RBP42 on slender bloodstream form *T. brucei* global proteome (a)** Schematic of quantitative proteomic (iTRAQ) experiment. Total cellular proteins from RBP42$^{Ty1}$ cells were extracted before (day 0) and after two days of RBP42 knockdown (day 2). Trypsin digested peptides from four replicate samples of each were labeled with iTRAQ tags (tag 117 – tag 121) and quantified by LC-MS/MS. **(b)** Box and whisker plots show similar distributions of normalized iTRAQ reporter ion intensities (log10) of ~5800 protein groups in all eight samples. Median values of all samples (x -axis) are also shown. **(c)** Immunoblot analysis of the total cellular proteins that were iTRAQ quantified. Anti-RBP42 antibody detects RBP42 protein. Clear depletions of RBP42 are evident in all four replicates on day 2. Anti-DRBD3 immunoblot is loading control. **(d)** Dot plot of RBP42 intensities (iTRAQ tags, log2) showing robust (> 5-fold) reduction of RBP42 protein. RBP42 protein levels from all eight samples are shown. **(e)** Correlation heatmap of iTRAQ reporter intensities shows two groups, before (day 0) and after RBP42 knockdown (day 2), indicating reproducibility of the analysis. Pearson correlation coefficients are shown.

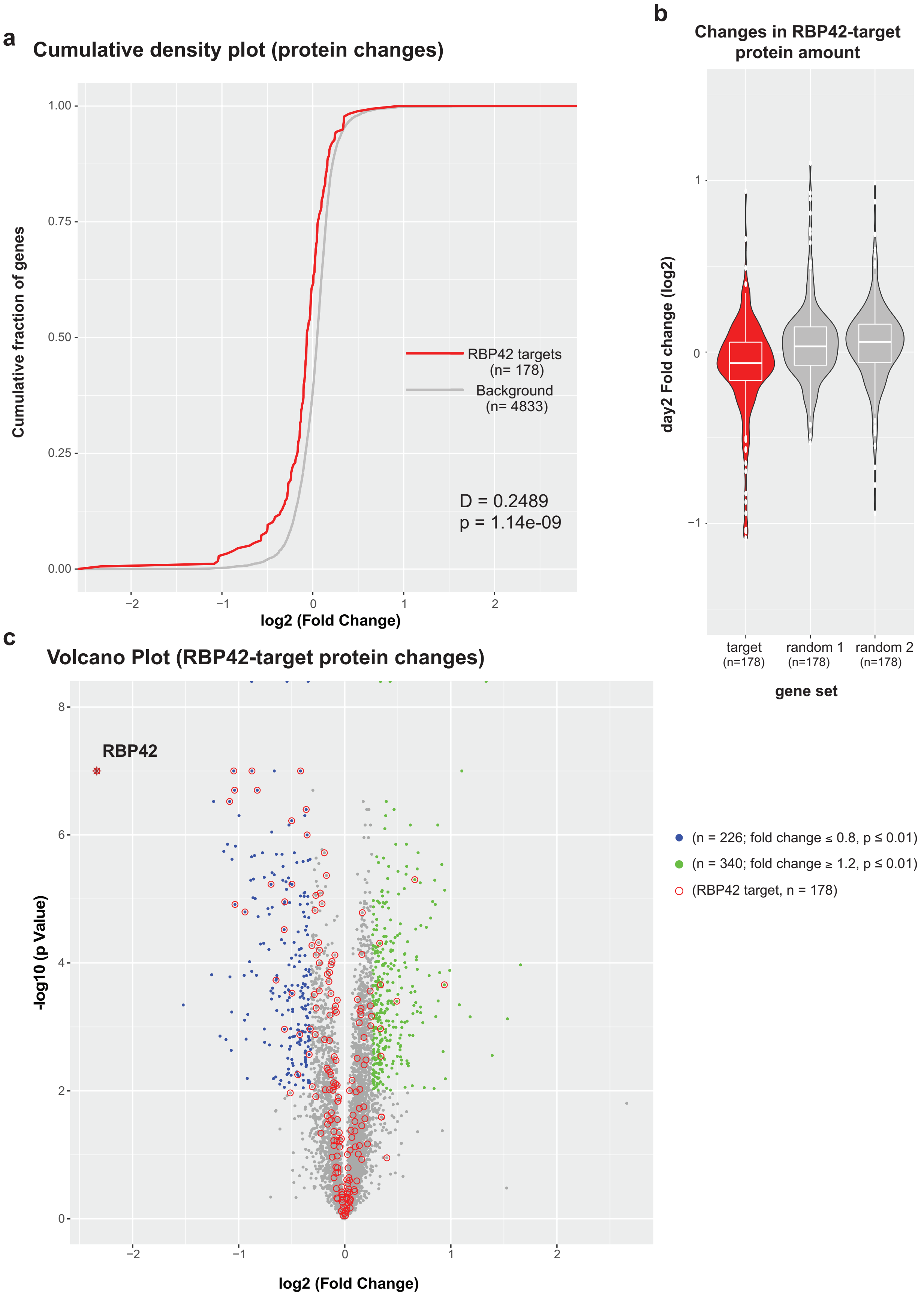
a
Cumulative density plot (protein changes)
1.00
0.75
0.50
0.25
0.00
Cumulative fraction of genes
−2
−1
0
1
2
log2 (Fold Change)
RBP42 targets (n= 178)
Background (n= 4833)
D = 0.2489
p = 1.14e-09
b
Changes in RBP42-target protein amount
1
0
−1
day2 Fold change (log2)
target (n=178)
random 1 (n=178)
random 2 (n=178)
gene set
c
Volcano Plot (RBP42-target protein changes)
8
6
4
2
0
-log10 (p Value)
RBP42
−2
−1
0
1
2
log2 (Fold Change)
(n = 226; fold change ≤ 0.8, p ≤ 0.01)
(n = 340; fold change ≥ 1.2, p ≤ 0.01)
(RBP42 target, n = 178)

**Fig S8. Loss of RBP42 causes reduction in target mRNA-encoded proteins**

**(a)** Cumulative distribution frequency (CDF) plot shows the empirical cumulative distribution of log2 fold changes for RBP42 target mRNA-encoded proteins (red) compared to non-target background proteins (grey) following RBP42 knockdown. The leftward shift of the RBP42-tagets (red) along the x-axis, compared to background (grey) indicates downregulation of the target proteins. The Kolmogorov-Smirnov (KS) test D-value and p-value are shown. **(b)** Violin plots showing the distribution of fold changes (log2) of the 178 RBP42 target mRNA-encoded proteins (target, red). Similar plots of two control protein sets of equal size, randomly sampled from the iTRAQ data, are also shown (random1 and random2, grey). Plot details are as in Fig S6. **(c)** Volcano plot, as in Figure S6, showing changes in protein levels (log2 fold changes) versus p-values (log10) for all (~5000) quantified protein (iTRAQ) after two days of RBP42 knockdown. The most significantly changed proteins (340 upregulated, green dots; 226 downregulated, blue dots), and the set of 178 RBP42 target-encoded proteins (red circle) are highlighted. All other proteins are shown in grey.

Volcano Plot (mRNA changes)

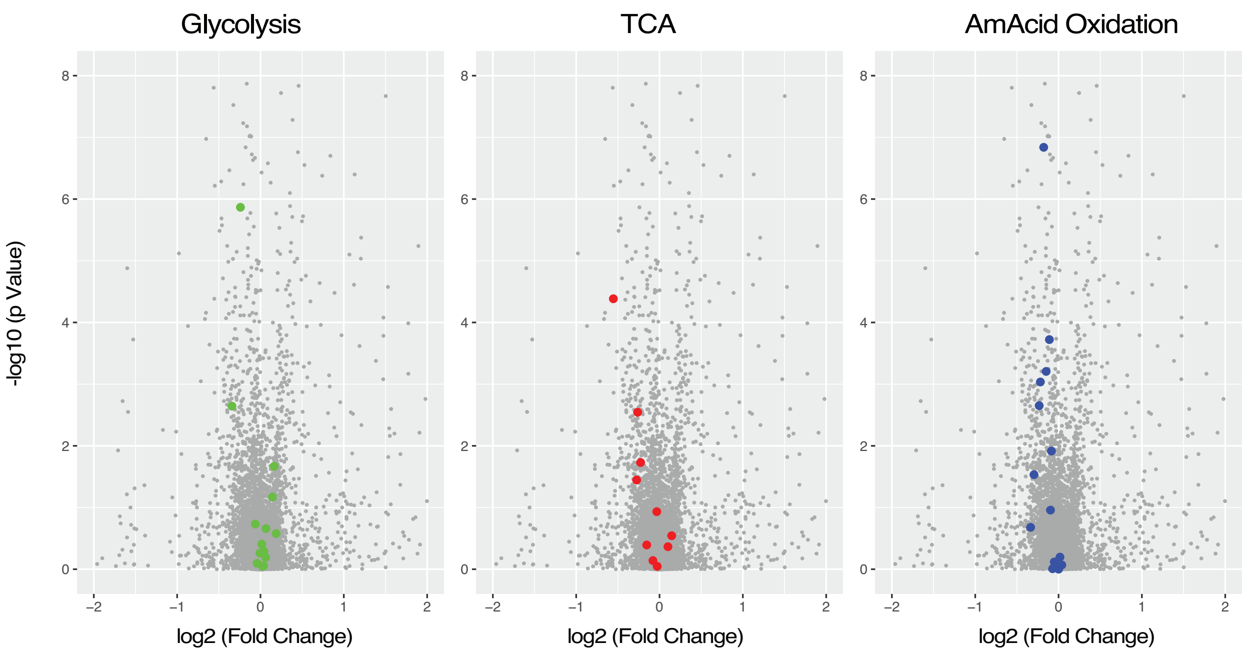


**Fig S9. Loss of RBP42 does not alter transcript levels**

Volcano plots, as in Figure 5, showing that loss of RBP42 does not alter mRNA levels of indicated gene cohorts, highlighted in color. mRNA levels ($log_2$-fold changes) versus significance p-values (-log10) for all protein coding genes, before and after two days of knockdown are shown.

**Glycolysis: protein and metabolite changes**

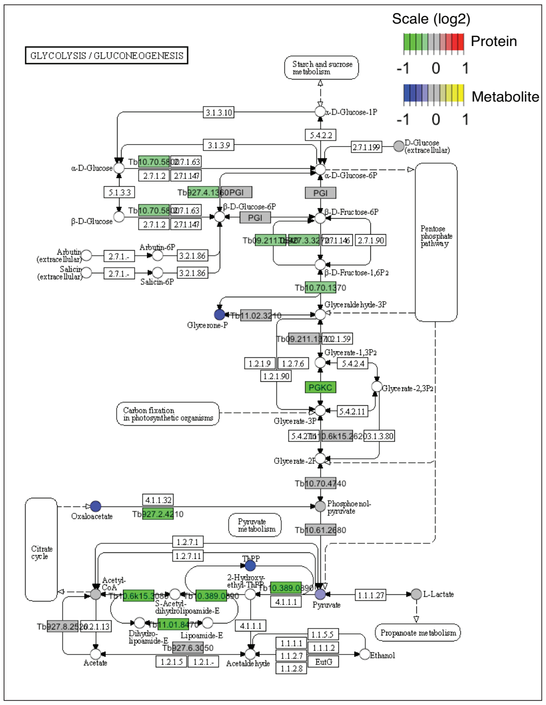


**TCA cycle: protein and metabolite changes**

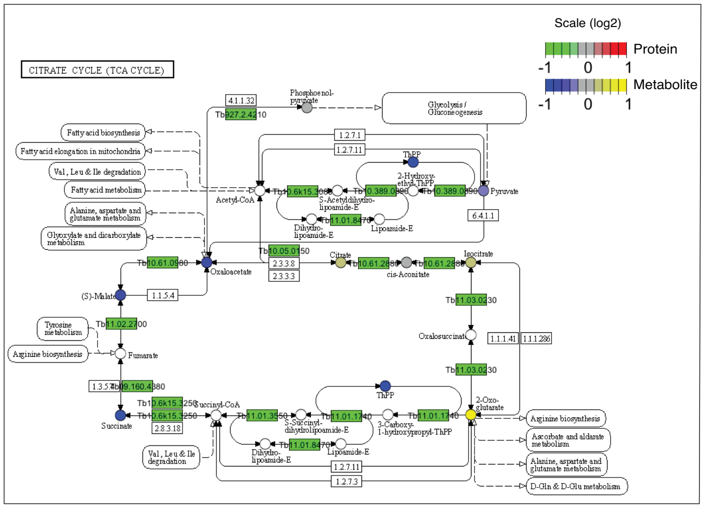


**Fig S10. Changes in proteins and metabolites following loss of RBP42**

KEGG map ( https://www.genome.jp/kegg/) view of *T. brucei* glycolysis and gluconeogenesis pathway (tbr00010) and TCA cycle pathway (tbr00020), rendered using Pathview (Luo, W.; Brouwer, C.: *Bioinformatics* **2013**, *29*, 1830–1831). Each box identifies a specific enzymatic step; each circle identifies a specific metabolic intermediate. Changes (log2 scale) in proteins and compounds are denoted using color ramps, green-to-red and blue-to-yellow respectively. No color implies that these enzymes and metabolites were not quantified. Note that the KEGG database identifies *T. brucei* genes using old ID.
